# The 22q11.2 region regulates presynaptic gene-products linked to schizophrenia

**DOI:** 10.1101/2021.09.22.461360

**Authors:** Ralda Nehme, Olli Pietiläinen, Mykyta Artomov, Matthew Tegtmeyer, Christina Bell, Andrea Ganna, Tarjinder Singh, Aditi Trehan, Vera Valakh, John Sherwood, Danielle Manning, Emily Peirent, Rhea Malik, Ellen J. Guss, Derek Hawes, Amanda Beccard, Anne M. Bara, Dane Z. Hazelbaker, Emanuela Zuccaro, Giulio Genovese, Alexander A Loboda, Anna Neumann, Christina Lilliehook, Outi Kuismin, Eija Hamalainen, Mitja Kurki, Christina M. Hultman, Anna K. Kähler, Joao A. Paulo, Jon Madison, Bruce Cohen, Donna McPhie, Rolf Adolfsson, Roy Perlis, Ricardo Dolmetsch, Samouil Farhi, Steven McCarroll, Steven Hyman, Ben Neale, Lindy E. Barrett, Wade Harper, Aarno Palotie, Mark Daly, Kevin Eggan

## Abstract

To study how the 22q11.2 deletion predisposes to psychiatric disease, we generated induced pluripotent stem cells from deletion carriers and controls, as well as utilized CRISPR/Cas9 to introduce the heterozygous deletion into a control cell line. Upon differentiation into neural progenitor cells, we found the deletion acted in trans to alter the abundance of transcripts associated with risk for neurodevelopmental disorders including Autism Spectrum Disorder. In more differentiated excitatory neurons, altered transcripts encoded presynaptic factors and were associated with genetic risk for schizophrenia, including common (per-SNP heritability p (*τ_c_*)= 4.2 x 10^-6^) and rare, loss of function variants (p = 1.29×10^-12^). These findings suggest a potential relationship between cellular states, developmental windows and susceptibility to psychiatric conditions with different ages of onset. To understand how the deletion contributed to these observed changes in gene expression, we developed and applied PPItools, which identifies the minimal protein-protein interaction network that best explains an observed set of gene expression alterations. We found that many of the genes in the 22q11.2 interval interact in presynaptic, proteasome, and JUN/FOS transcriptional pathways that underlie the broader alterations in psychiatric risk gene expression we identified. Our findings suggest that the 22q11.2 deletion impacts genes and pathways that may converge with risk loci implicated by psychiatric genetic studies to influence disease manifestation in each deletion carrier.

## Introduction

Heterozygous deletions of the 22q11.2 chromosomal interval occur approximately once in every 4,000 live births^1^. This deletion confers a risk of developing several symptomatically diverse neuropsychiatric conditions including intellectual disability (ID), Autism Spectrum Disorder (ASD) and schizophrenia^2–7^. In fact, deletion of 22q11.2 confers the largest effect of any known genetic risk factor for schizophrenia^8^.

Unlike the 22q13.3 deletion syndrome, where risk of mental illness can largely be explained by reduced function of a single gene (*SHANK3*)^9^, mutations in no one gene within the 22q11.2 deletion can explain the predisposition for psychiatric disease it confers. As a result, the pathways through which the 22q11.2 deletion contributes to ASD and schizophrenia risk remain poorly understood. Mouse models have served as an initial in road for identifying genes within the deletion that function in brain development and behavior. Overall, studies with rodent models suggest that several genes in the syntenic chromosomal interval including *Dgcr8*, *Ranbp1, Rtn4r,* and *Zdhhc8* have important nervous system functions^10–21^. However, imperfect alignment between mouse behavioral phenotypes and psychiatric symptoms have left uncertainty concerning which, or how many of their human orthologs play a role in mental illness.

More recent studies now suggest that the genetic background of 22q11.2 deletion carriers contributes meaningfully to their likelihood of developing one psychiatric condition or another. For instance, deletion carriers that also harbor an additional copy number variant (CNV) elsewhere in the genome displayed a higher risk of developing schizophrenia^22^. Additionally, analysis of polygenic risk scores calculated using data from genome wide association studies (GWAS) suggests that an increased burden of common risk variants can act in concert with the 22q11.2 deletion to further increase overall risk for psychosis^23–25^. These observations clearly indicate the 22q11.2 deletion can at least act together with alterations in genetic pathways affected by additional risk variants. This raises the possibility that the deletion may converge on disease mechanisms that act in both ASD and schizophrenia.

We reasoned that finding the points of convergence between the effects of the 22q11.2 deletion and other human genetic variants implicated in psychiatric disorders could provide a view into which genes present in the deletion, or pathways altered by it, contribute to mental illness. To identify such intersections, we opted to examine transcriptional changes in multiple stages of excitatory neuronal differentiation, given that genetic studies of ASD and schizophrenia have implicated genes that act during neuronal development and differentiation^26–29^, and in neuronal processes including excitatory transmission^30–32^. We therefore carried out RNA sequencing at three distinct stages of excitatory neuronal differentiation using induced pluripotent stem cells (iPSCs) from 22q11.2 carriers and non-carrier controls. In order to establish a causal link between the deletion and the transcriptional effects we also utilized gene editing to delete the chromosomal region in a control cell line. To robustly induce neuronal differentiation, we utilized an approach we previously described where Ngn2 expression^33^ is coupled with forebrain patterning to produce homogenous populations of excitatory neurons with features similar to those found in the superficial layers of the early cortex^34^. We have previously characterized the cells generated by this approach using immunostaining, qPCR, single-cell RNA sequencing, whole-cell patch clamp, multi-electrode arrays and optical electrophysiology, and demonstrated reproducibility across multiple cell lines^34–37^.

Over the course of excitatory neuronal differentiation, we found that the 22q11.2 deletion acted in trans to significantly alter the expression of many genes with established genetic associations with neurodevelopmental disorders in progenitors, and schizophrenia in differentiated neurons. To ask, in an unbiased manner, which pathways and genes were likely responsible for these changes, we developed an approach for identifying protein-protein interaction (PPI) networks that best explain a particular change in gene expression. This method, called PPItools, suggested that the 22q11.2 interval regulates the expression of genes in proliferative, presynaptic, proteasomal and JUN/FOS pathways. Finally, we found that cell lines with isogenic deletion of 22q11.2 recapitulated most of the changes observed in the patient-based cohort, including increased levels of the *MEF2C* transcription factor in neuronal progenitor cells and decreased expression of presynaptic proteins such as SV2A and NRXN1 in neurons.

## Results

### Pilot study and power calculations

The 22q11.2 deletion syndrome is associated with a wide spectrum of psychiatric conditions, which differ from person to person, and by age of diagnosis. To study the effects of the deletion, we both collected and derived hiPSC lines from patient carriers as well as non-carrier controls (Fig.1a-f, Extended Data Fig. 1a and Extended Data Table 1).

**Fig. 1.**
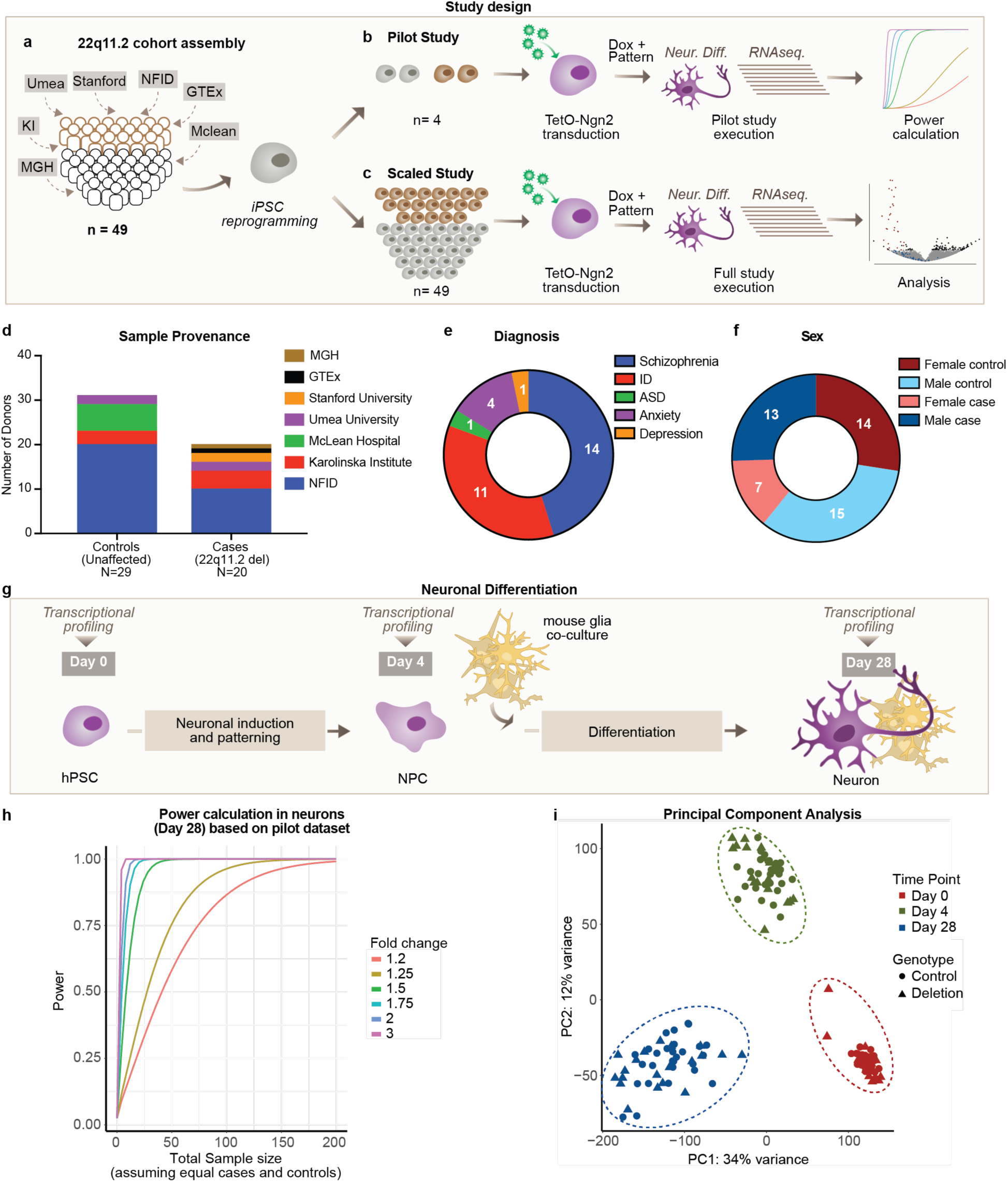
Design of a statistically powered study to determine the impact of 22q11.2 deletion on gene expression. **a**, Final sample set composed of 20 cell lines with 22q11.2 deletion (brown) and 29 controls (grey), collected at seven locations (MGH: Massachusetts General Hospital, KI: Karolinska Institute, Umea: Umeå University, NFID: Northern Finnish Intellectual Disability Cohort (Institute for Molecular Medicine Finland), GTEx: Genotype-Tissue Expression Project, Mclean: Mclean Hospital) **b**, Pilot study using four hiPSC lines differentiated into neurons through transduction with TetO-Ngn2, Ub-rtTA and TetO-GFP lentivirus and subjected to RNA sequencing. RNA abundances were then used to estimate the appropriate sample size for differential gene expression for the final study. **c**, The final dataset consisted of 49 cell lines that were differentiated and subjected to RNA sequencing. **d**, Provenance **e**, Diagnosis and **f**, Sex of the samples in the final cohort. **g**, Neuronal differentiation protocol, previously published and characterized^34,37^ consisting of the combination of Ngn2 overexpression with forebrain patterning using small molecules (SB431542, LDN193189 and XAV939). Samples were harvested for RNA sequencing at the stem cell (day 0), neuronal progenitor cell (NPCs) (day 4) and neuronal (day 28) stages. **h**, Power estimation in the pilot dataset for median expressed genes (24 read counts) for different fold-changes and sample sizes in neurons. **i**, Principal component analysis (PCA) of RNA sequencing data from the full study.

To estimate the sample size needed to be powered to detect gene expression changes, we performed a pilot study with two control and two 22q11.2 deletion iPSC lines, each from a distinct donor. The presence of the 3Mb 22q11.2 deletion provided an internal control with a built-in expectation for a set of known deleted genes and their anticipated magnitude of change. Thus, we reasoned this small study would allow us to detect the 50% reduction in the abundance of transcripts originating from within the deletion as well as changes in expression of genes outside of the deletion that were of a similar magnitude. We induced neuronal differentiation using a published, well-characterized approach, combining the overexpression of Ngn2 with small molecule patterning^37^ (Fig. 1g) , and completed RNA-sequencing at three cellular stages: human pluripotent stem cells (hPSCs, day 0 of differentiation), neuronal progenitor-like cells (NPCs, day 4 of differentiation)^37^, and in functional excitatory neurons displaying synaptic connectivity ^34^ (day 28 of differentiation) (Extended Data Table 1, Extended Data Fig. 1b-e). Following RNA-sequencing, we mapped reads to the Ensembl human genome assembly (GRCh37/hg19). We detected one or more reads for 51 protein coding genes that mapped to the 22q11.2 deletion region, in the four lines at any one differentiation stage. On one hand, we were reassured to observe a systematic reduction in the abundances of RNAs encoded by genes mapping in the deletion, with the majority exhibiting fold-changes between -1.5 and -2 in deletion cells relative to controls. On the other hand, this decrease in RNA levels was indistinguishable from sample-to-sample variance on an individual gene level (after correcting for multiple testing), underscoring the limitations of a small sample size (Extended Data Fig. 1c-e). Only when we considered reads from the genes in the deleted region in aggregate could we observe a statistically significant reduction in gene expression between the deletion carriers and controls (p(hPSCs) = 4.13×10^-19^, p(NPCs) = 1.58×10^-18^, and p(neurons) = 2.93×10^-15^, Mann-Whitney test).

Using our pilot sequencing data, we estimated that for genes expressed above the median, a sample size of > 20 carrier and > 20 control iPSC lines would yield on average >80% power to detect fold-changes of 1.35 across each of the three cell stages (Fig. 1h, Extended Data Fig. 1e,f).

### Profiling an expanded 22q11.2 cohort

Guided by our power calculations, we assembled a collection of 20 (7 female, 13 male) 22q11.2 deletion carrier and 29 (14 female, 15 male) control iPSC lines, each derived from a distinct individual. It has been found that the size of the deletion doesn’t seem to correlate with diagnosis or severity of the conditions, as patients with either the most common 3Mb deletion or smaller nested deletions appear to have similar diagnoses^3,7,38,39^. We thus decided to examine the impact of the deletion (agnostic to size or diagnosis) on gene expression during neuronal development. We performed RNA sequencing in hPSCs, NPCs and excitatory neurons for each of the 49 cell lines (in triplicates, N=441 total RNA sequencing libraries in mixed pools of both genotypes to minimize technical biases). With these data in hand, we revisited our initial power estimates and found that in the larger data set we achieved over 80% power to detect fold changes ≥1.5 of all detected protein coding genes (Extended Data Fig. 1h) across developmental stages.

Consistent with previous findings using the same neuronal differentiation approach^34,37^, differentiation down a neuronal trajectory resulted in a global change of gene expression between each cellular stage analyzed (day 0 iPSC, day 4 NPC, day 28 excitatory neuron). Principal component analysis (PCA) indicated that the primary component of variation between the samples was days of neuronal differentiation (PC1+2 = 46% of variance) (Fig. 1i, Extended Data Fig. 1g). We found close clustering of the samples from the 49 lines within a given differentiation time point within PC1 and PC2, suggesting a reproducible and reliable differentiation had occurred across the entirety of our experiments (Fig. 1g). This conclusion was supported by joint analysis of the data indicating that across the 49 cell lines, 4 pluripotency associated genes were robustly expressed at day 0 and then rapidly silenced, while 7 representative NPC genes became expressed at day 4 with the strong emergence of 7 prototypical neuronal genes at day 28 (Fig. 2a, Extended Data Fig. 2c).

**Fig. 2.**
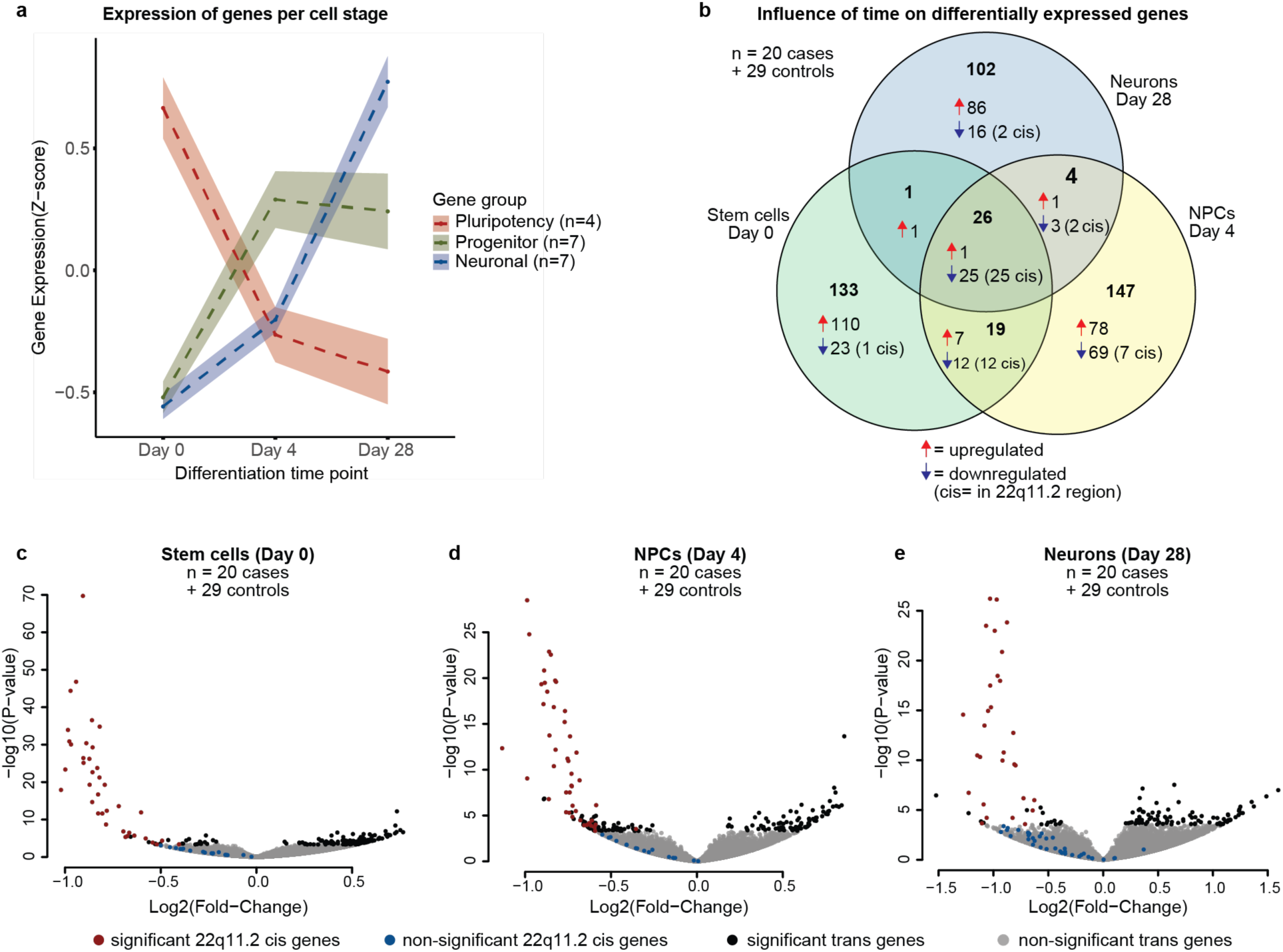
Cell-type specific effects of the 22q11.2 deletion. **a**, Expression of selected marker genes for defined specific cell stages by suppression of genes related to pluripotency (*SOX2*, *OCT4*, *NANOG*, *MKI67*) and up-regulation of genes characteristic for neural progenitor cells (*NEUROD1*, *SOX2*, *EMX2*, *OTX2*, *HES1*, *MSI1*, *MKI67*) and mature neurons (*NEUN*, *SYN1*, *DCX*, *MAP2*, *TUJ1*, *NCAM*, *MAPT*) as the differentiation progresses (gene lists also provided in Extended Data Fig. 2c). **b**, Venn Diagramm highlighting the number and directionality of shared and unique differentially expressed genes between deletion carriers and controls at each cell stage. Genes within the deletion region (cis) are mostly shared across development stages, whereas genes outside the deletion region (trans) are cell-stage specific. **c-e**, Volcano plots showing differential gene expression in stem cells (c), NPCs (d) and neurons (e) Significantly differentially expressed genes (FDR<5%) within the deletion region are presented in red and outside deletion in black. Non-significant genes in deletion region are presented in blue.

### 22q11.2 effects on transcript abundance

We next proceeded to ask the important question of how the 22q11.2 deletion status influenced gene expression during neuronal differentiation and first considered genes within the deletion. We observed a nominally significant reduction in RNA levels for 51 protein coding genes in the deletion region (p<0.05, red and blue dots, Fig. 2c-e) with 49 of these transcripts yielding significantly reduced abundance in at least one time point (FDR <0.05) and 25 significantly reduced in all 3 stages (FDR <0.05) (Fig. 2b, Extended Data Fig. 2a). These findings in excitatory neuronal cells were in line with previous reports using either mixed monolayer cultures of inhibitory and excitatory neurons carrying the 22q11.2 deletion^40^ or organoids consisting of multiple cell types including glutamatergic neurons and astrocytes^41^. We found that for genes mapping to the deletion, which showed a significant change in their expression, the deletion genotype explained, on average, 42 to 52% of all variance in their expression (Extended data Fig. 3a-d). Included in these 49 significantly less abundant transcripts originating from the 22q11.2 locus, were seven that are highly intolerant for loss of function variants as measured by pLI score^42^, which ranks genes from most tolerant (pLI=0) to most intolerant (pLI=1). These seven genes that have a pLI score > 0.9 (*UF1DL, HIRA, DGCR8, ZDHHC8, MED15, TBX1*) have been previously suggested to play role in some of the congenital phenotypes associated with the deletion in other tissues^43^. Together, our analyses indicate that our transcriptional phenotyping was sufficiently sensitive to allow the successful detection of the 50% decrease in expression of the hemizygote genes found in the deletion region.

### Cell-type specific effects of 22q11.2 deletion

After validating our ability to detect the altered expression of many genes within the deletion, we next explored differentially expressed transcripts originating from loci outside of the deletion. In fact, the majority (89%) of the genes differentially expressed in 22q11.2 carrier’s cells were located outside the deletion region (n=383 genes) (Fig. 2b). In total, the trans effects of the deletion explained on average 18% of the total variance in gene expression across all data sets (Extended data Fig. 3a-d). Plotting the test statistic from the differential expression for every gene relative to its position in the genome suggested that there was no major positional clustering of differentially regulated genes to specific chromosomal regions outside the deletion area (Extended data Fig. 3e, day 28 example). Notably, only one gene, *CAB39L* on chromosome 13, was significantly induced in carriers at all stages (Extended Data Fig. 2b). Upon reviewing published data sets, we found that *CAB39L* expression was also induced in blood cells isolated from 22q11.2 deletion carriers^44^, suggesting that upregulation of this gene is likely to be associated with the 22q11.2 deletion in many cell types.

While genes within the 22q11.2 deletion region were regulated in the same direction at all developmental stages, the set of differentially expressed genes outside the deletion region was different for each stage. In contrast to the conserved downregulation of genes within 22q11.2 across the three distinct time points we analyzed, except for *CAB39L*, the specific identity of the remaining differentially expressed genes was distinct at each differentiation stage assessed (372 cell stage-specific genes). Importantly, the apparently discontinuous effects of the deletion between the cell stages were not the trivial result of certain transcripts failing to be detected because of barely falling outside a certain significance threshold. That is, in controls, the affected genes were expressed in all cell stages with little change in their overall average RNA abundance between stages, ensuring reliable detection across all stages (Table S4 and Extended Data Fig. 4a,b). As a result, fold-changes in “trans” genes between carriers and controls were only modestly correlated between NPCs and hPSCs (ρ=0.28, p=3 × 10^-8^) and NPCs and neurons (ρ=0.23, p=3 × 10^-6^), while no correlation was observed between fold changes in hPSCs and neurons (ρ=0.06, p=0.25). Overall, these findings suggest that the 22q11.2 deletion has a temporally-dependent influence on gene expression, altering the abundance of distinct sets of transcripts as neuronal differentiation unfolds.

Lastly, our cohort included 18 cell lines with full length 22q11.2 deletion, along with two cell lines with nested 22q11.2 deletion. To verify that the cell lines with shorter deletion did not result in a different transcriptional signature, we repeated the differential gene expression analysis in day 28 neurons without the lines with short, nested deletions (SCBB-1430 and SCBB-1961, Table 1). We found that the differences in gene expression between the remaining deletion carriers and controls correlated strongly with those obtained in the complete data set (r=0.92 for genes with adjusted p-value<0.05), suggesting that the observed gene expression differences were robust also in the presence of the shorter deletions.

### Transcript alterations in hPSCs and NPCs

The phenotypes that are found in a subset of 22q11.2 deletion carriers during early childhood^3^ led us to ask if the genes we identified to be differentially expressed in deletion carriers at initial differentiation stages (hPSCs and NPCs) were genetically associated with neurodevelopmental disorders, including autism and intellectual disability. We included likely disease-causing genes from the Deciphering Developmental Delay (DDD) project, and a recent, large exome-sequencing study in autism (n=295 total neurodevelopmental disorders, NDD, genes)^27,45,46^ (Table S5). Of the 432 genes we found differentially expressed in deletion carriers, 10 were NDD genes (hPSCs: *FOXG1, ELAVL3;* NPCs: *PAX6*, *MEF2C*, *FOXP2*, *NR2F1*, *MAF*, *PAX5;* neurons: *KMT2C*, *MKX;* OR = 1.85, p=0.046 (for all 432 genes) (Tables S1-S3). We took particular note of *MEF2C* as it is also implicated in schizophrenia through GWAS^47^ and is known to encode a transcriptional regulator that participates in activity-dependent regulation of immediate early genes such as *JUN* and *FOS* ^48^. *MEF2C* has been shown to be repressed by the transcription factor *TBX1*, which is encoded by a gene within the 22q11.2 interval^49,50^.

Proteins encoded by genes harboring causal mutations for a particular phenotype in Mendelian disorders have been shown to have more physical connections between one another than unrelated proteins^51^. We therefore wondered whether the transcripts expressed from within the 22q11.2 deletion and the transcripts with altered abundance in trans in deletion carriers encoded proteins that together had more than the expected number of interactions with proteins originating from loci genetically linked with NDD. As this is a question of broader relevance for connecting protein interaction data, changes in gene expression, and genetic data, we wrote a software package (PPItools, https://github.com/alexloboda/PPItools) to enable this analysis.

In this instance we used PPItools to identify the protein-protein interactions (PPI) from the InWeb database^52^ of the differentially expressed gene products that we identified at each stage of neural differentiation and analyzed them for an apparent excess of genes implicated in NDD in this network. We used a curated list of NDD genes that comprised 295 genes that have been previously reported to have excess of deleterious variants in patients with ASD, and ID ^45,46,53^ (Table S5). To ask whether this enrichment for NDD implicated interacting proteins was likely to have occurred by chance, we performed 1000 random permutations of sets of expressed proteins of the same size while constraining the scale and complexity of the network. These analyses confirmed that genes we found to be differentially expressed early in differentiation (in hPSCs and NPCs) were significantly more likely to interact with gene products associated with NDD (p<0.001, Extended Data Fig. 4c). While there remained a modest enrichment for differentially expressed genes in excitatory neurons for interaction with NDD gene products, this enrichment was not significant.

To further control our observation, we asked whether the protein interaction network we identified at each time point showed any enrichment for genes linked with an unrelated condition, inflammatory bowel disease (IBD), or with a neurological condition, Parkinson’s Disease (PD). As expected, there were no significant enrichments for IBD related gene products within the protein interaction networks identified at any of the differentiation time points analyzed, and no enrichment for PD related gene products in NPCs or neurons (Extended Data Fig. 4c, Table S5). Thus, our results demonstrate that within hPSCs and NPCs, there is indeed a convergence between genes within the 22q11.2 deletion and the transcripts altered in trans by the deletion with genes products that when mutated cause human neurodevelopmental disorders.

### Schizophrenia heritability enrichment in neurons

Given that we had found an initial convergence between the effects of the 22q11.2 deletion and the abundance of certain transcripts linked through rare variant analyses to NDD as well as with a broader collection of PPI networks implicated in NDD, we next proceeded to ask whether the transcripts that we had found to be altered in deletion carrier cells were enriched for additional genetic signals in mental illness. To investigate this possibility, we utilized the genes we identified to have significantly altered expression (FDR < 0.05) in differentiating cells from 22q11.2 carriers as a substrate for linkage disequilibrium (LD)-score regression^30^. For this analysis we used GWAS summary statistics from the psychiatric genomics consortium (PGC), as well as educational attainment studies^54–59^ to ask whether variants in 22q11.2-differentially expressed genes and their surrounding genomic regions contribute disproportionately to the polygenic heritability of five neuropsychiatric disorders (schizophrenia, bipolar disorder, major depressive disorder, autism spectrum disorder, and ADHD) . We applied two statistics to estimate heritability enrichment in LD-score regression: per-SNP heritability and total heritability enrichment. We found suggestive evidence for a modest increase in per-SNP heritability for schizophrenia among genes differentially expressed in neurons *τ_c_*=6.1 x 10^-8^; p=0.0088 for 196 genes, FDR <5% and *τ_c_*=1.5 x 10^-8^; p= 0.01 after examining all 4,192 genes with nominally significant differences in expression, p<0.05, 2,864 up genes and 1,328 down genes, respectively) (Fig.3a). Analysis of up- and down-regulated genes (p<0.05) separately revealed that the increase in the per-SNP heritability was accounted for by transcripts that were more abundant in deletion carrier neurons (p (*τ_c_*)= 4.2 x 10^-6^ p(Bonferroni)=0.0003) (Fig.3a, Table S6). Our findings were unlikely to be the result of neurons merely expressing increased levels of genes relevant for these psychiatric conditions: permutation with 100 random gene lists produced from our neuronal data and matched for expression level, indicated that the per-SNP heritability enrichment in genes we found to be induced in deletion carrier neurons was ∼10,000-times more significant than any random gene set (Extended Data Fig.5a-c). We found a similar trend when examining the total heritability accounted for by variants in these genes, where we found an increase in heritability enrichment for bipolar disorder and educational attainment in addition to schizophrenia (Extended Data Fig. 5d,e, Table S6).

**Fig. 3.**
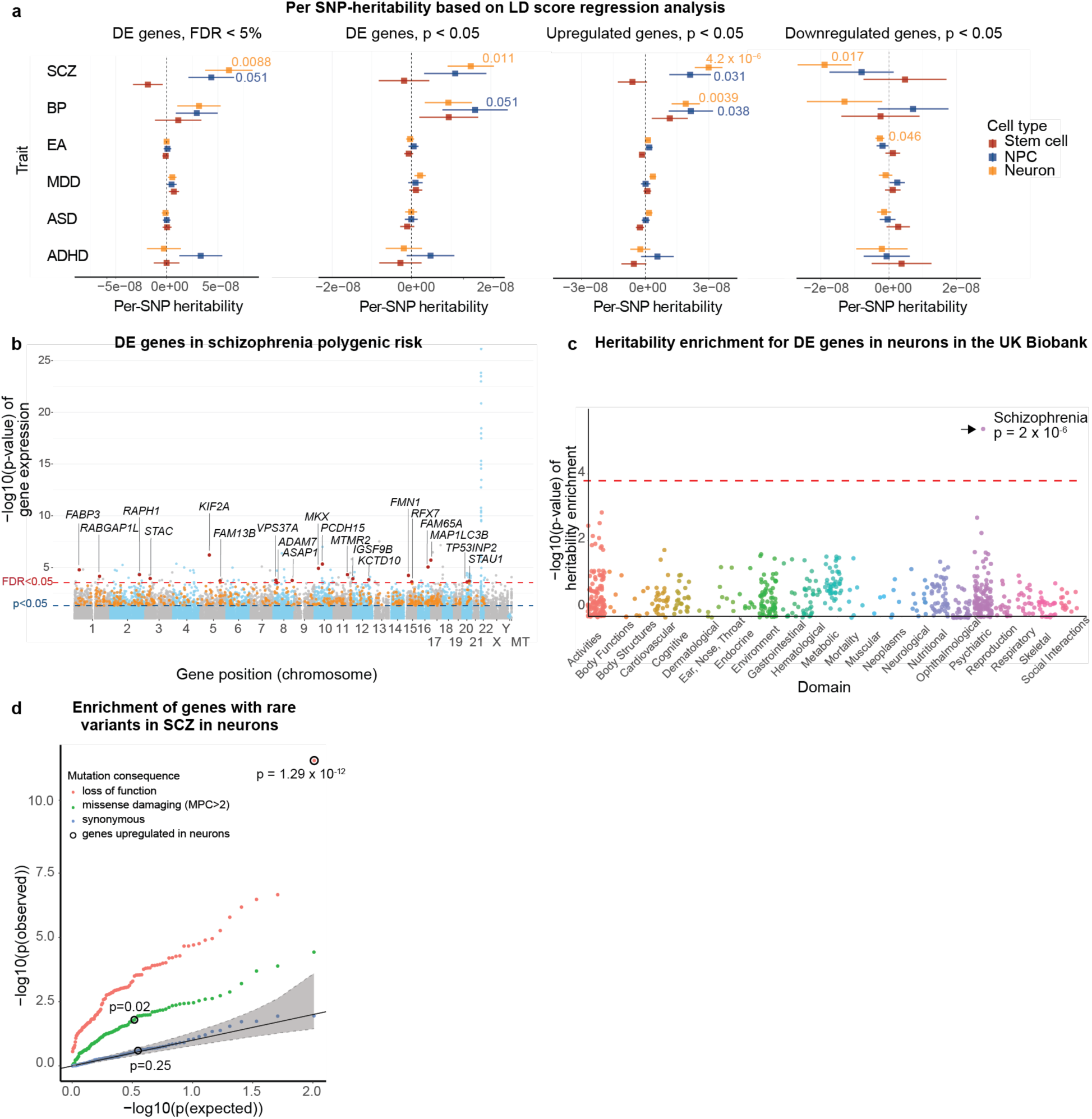
Heritability enrichment for schizophrenia risk genes 22q11.2 deletion neurons. **a**, Marginal enrichment in per-SNP heritability explained by common (MAF > 5%) variants within 100kb of genes differentially expressed, estimated by LD Score regression. Six traits were analyzed: SCZ=schizophrenia, BP=bipolar disorder, EA=educational attainment, MDD=major depressive disorder, ASD=autism spectrum disorder, ADHD=attention deficit hyperactivity disorder, at all three cell stages, showing enrichment for schizophrenia most prominently in genes upregulated in 22q11.2 deletion neurons. DE=differentially expressed. Four groups of DE genes were analyzed. Right, all DE genes with an FDR < 5%. Middle right, all nominally significant DE genes (p<0.05). Middle left, all nominally significant upregulated DE genes (p<0.05). Left, all nominally significant downregulated DE genes (p<0.05). **b**, DE genes in 22q11.2 neurons with nominally significant gene-wise association to schizophrenia from MAGMA (pg<0.05). **c**, GWAS summary statistics for 650 traits from the UK-biobank showing significant enrichment for heritability only for schizophrenia (p=2×10^-6^) in genes upregulated in deletion neurons. **d**, qq plot of p-values for the enrichment of rare coding LoF, missense damaging or synonymous variants in schizophrenia patients in genes upregulated in deletion neurons (circled in black) and 100 random gene sets matched by expression level to the upregulated genes.

To again query the relationship between differentially expressed genes in 22q11.2 deletion neurons and common genetic variants more broadly associated with psychiatric illness, but with a different set of statistical assumptions, we applied multiple-regression for competitive gene-set analysis in MAGMA-software^60^. Like results from the LD-score regression analysis, genes whose transcripts were more abundant in 22q11.2 deletion neurons were more strongly associated with schizophrenia than the rest of the genome (p=5.6 x 10^-7^, p(Bonferroni)=4.03 x 10^-5^) (Extended Data Fig. 6a). Altogether, 20 genes with nominally significant gene-wise association to schizophrenia from MAGMA (p_g_<0.05) were significantly differentially expressed in deletion neurons (Fig. 3b, Extended Data Fig. 6c). Repeating the analysis with 100 random gene lists generated from our expression data confirmed that this result was unlikely to have arisen merely as a result of examining these neuronal cells (Extended Data Fig. 6b).

To determine if this association between 22q11.2 deletion induced genes and schizophrenia heritability was replicable and to determine the specificity of this signal, we used summary statistics from an independent GWAS dataset of 650 heritable traits from the UK-biobank. Strikingly, LD-score regression showed the genes upregulated in 22q11.2 deletion neurons harbored significant heritability enrichment for schizophrenia, but not for the other traits (Fig.3c). Overall, our findings indicated that excitatory neurons harboring the 22q11.2 deletion exhibited increased abundance of transcripts from genes that underlie schizophrenia heritability, but that the deletion did not have such a detectable effect at earlier stages of differentiation.

### Schizophrenia rare variant enrichment in neurons

Exome sequencing at increasing scale has begun to reveal a burden of rare protein damaging variants in schizophrenia patients, complementing the genetic signal of common regulatory variants emerging from GWAS^61–63^. In contrast to the common variant polygenic risk, which arises incrementally from many small-effect variants, the schizophrenia-associated rare variants identified so far act with strong individual effects. While there is evidence for common and rare risk variants in schizophrenia mapping to shared chromosomal intervals^32^, so far the two forms of variation implicate largely distinct sets of genes. We therefore asked whether the 22q11.2 deletion also effects the expression of genes that harbor rare coding variants, identified by the schizophrenia exome meta-analysis consortium (SCHEMA) in schizophrenia patients^63–65^. We initially focused on genes upregulated in neurons from 22q11.2 carriers (n=2,864 genes at p < 0.05) and used 100 random gene lists matched for their expression levels in our excitatory neurons as controls. This analysis revealed two interesting results: First, alterations in the expression of genes harboring a burden of loss of function mutations in schizophrenia were significantly enriched in excitatory neurons from 22q11.2 carriers (Fig. 3d red dots, 57/100 random gene lists assessed p <0.05). Second, that the 2,864 transcripts within these neurons whose expression were increased in 22q11.2 deletion carriers were substantially more significantly enriched for loss of function variants than any of the random gene lists we sampled (Fig. 3d, red dot black circle; p = 1.29×10^-12^). This enrichment signal was substantially reduced for missense mutations in schizophrenia patients and absent for synonymous variants (green and blue dots Fig. 3d). We further examined the differential expression results for genes with significant burden of deleterious mutations in schizophrenia patients in SCHEMA. One transcript encoded by *ZMYM2* out of 32 significant genes from SCHEMA was significantly changed in the deletion lines (FDR<5%). Seven additional genes (*RB1CC1*, *AKAP11*, *ASH1L*, *GRIA3*, *SV2A*, *PCLO*, *DNM3*) were nominally significantly changed in the deletion neurons. Remarkably, all eight SCHEMA genes were upregulated in the deletion carrier neurons.

Consistent with the notion that we were analyzing a disease relevant cell type, our rare-variant burden analyses indicated that the excitatory neurons we produced from both cases and controls expressed a significant excess of genes harboring rare pathogenic coding variants in schizophrenia patients. However, our analysis further indicated that in excitatory neurons the 22q11.2 deletion was specifically associated with alterations in a set of genes that were even more markedly enriched for rare loss of function variants in schizophrenia patients (Fig. 3d, circled dot). Like our common variants analyses, genes whose expression we found altered in pluripotent stem cells and NPCs harboring the 22q11.2 deletion did not exhibit this excess of rare coding variants schizophrenia (Extended data Fig. 7a, Table S7).

### Protein-protein interaction networks associated with transcriptional changes

As the number of trans acting effects of the deletion on transcripts linked to psychiatric illness were substantial, we sought an unbiased approach for identifying the pathways that could be contributing to their alterations. Ideally, such a method would also have the capacity to identify potential connections to gene products originating from within the deletion interval itself. To this end, we used PPI data^52^ to search for the smallest number of biochemical interactions that could explain the most prominent transcriptional changes in deletion carriers. To facilitate this effort, we developed a new tool (included in the R-package “PPItools”, see methods) that scores observed p-values from differential expression to construct a node weighted graph with the strongest cumulative association with the deletion genotype at each cell stage (most-weighted connected subgraph, MWCS). We then performed 1000 permutations on p-values from differential expression while preserving the node degrees, to ensure that the connected gene-products were unlikely to occur in the subgraph by chance alone (p<0.05, Table S8) (Extended Data Fig. 8a). This analysis revealed that the minimal interaction networks for each of the three stages of differentiation were predominantly composed of proteins encoded by genes located within the 22q11.2 deletion, that were in turn interconnected with proteins encoded by genes residing outside of the deletion (Extended Data Fig. 8b,c and Fig. 4g).

**Fig. 4.**
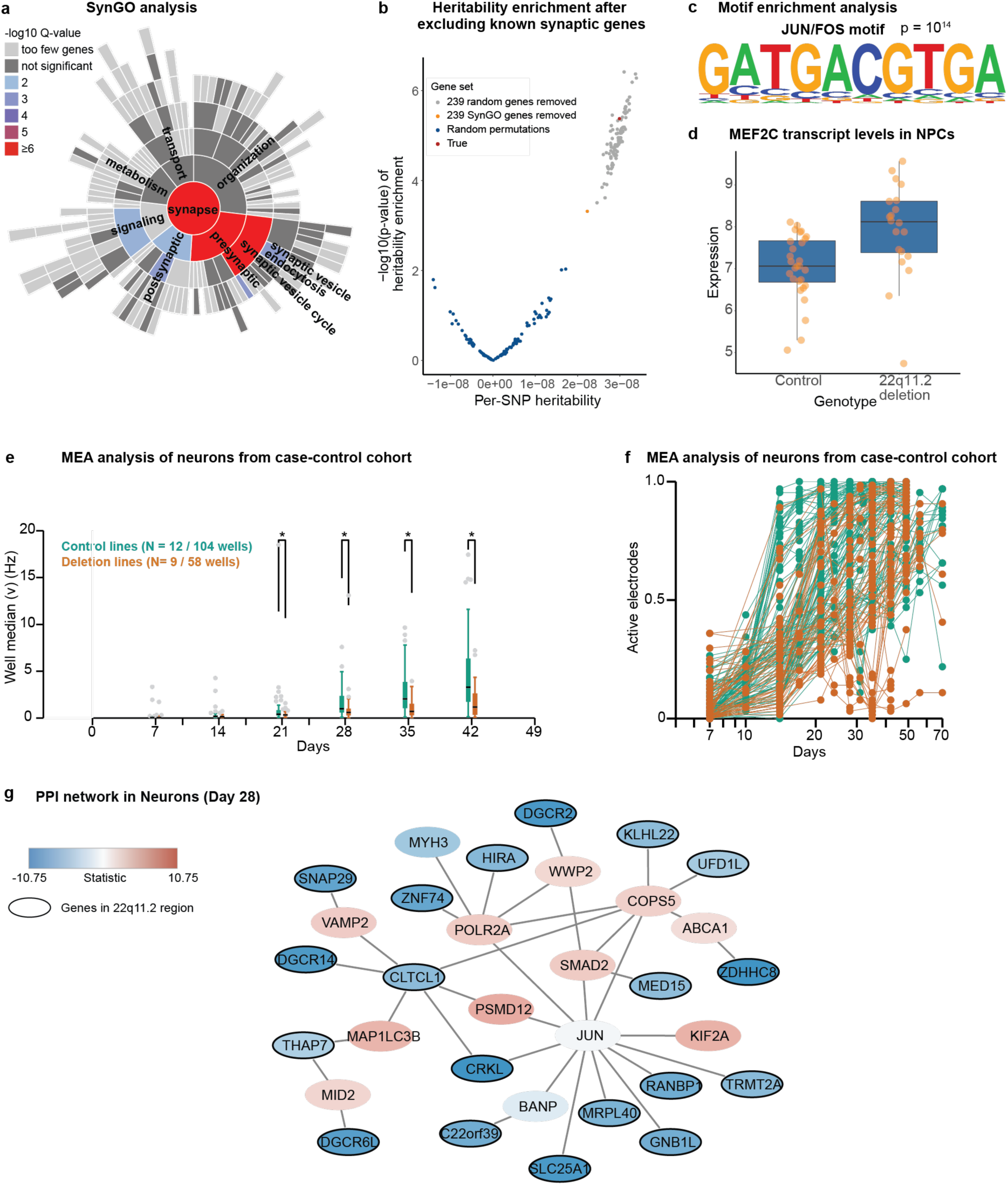
Impact of the 22q11.2 deletion on synaptic gene expression and network activity. **a**, SynGO annotation for genes upregulated in neurons showing enrichment for synaptic processes. **b**, Heritability enrichment for schizophrenia after excluding the 239 genes with SynGO annotation. **c**, Motif Enrichment analysis in upregulated genes (p <0.05), showing enrichment of JUN / FOS targets. **d**, *MEF2C* is upregulated in NPC of 22q11.2 deletion carriers. **e,** Spike count (mean number of spikes in a 10 second period). The activity of neurons derived from control (green, N = 12 lines, 104 wells) is compared to neurons from cases with 22q11.2 deletion (N = 9 lines, 54 wells). **f,** Proportion of electrodes detecting spontaneous activity, against the number of days post-induction. **g,** The most weighted sub-cluster graph for protein-protein interactions (PPI) for differentially expressed genes in neurons.

In pluripotent stem cells, we found that the most weighted subgraph contained 53 node proteins, 26 of which were encoded by genes mapping to the deletion (Extended Data Fig. 8b). These nodes were organized around several hub proteins encoded by genes that map outside the deletion. These included MYC, p53 (TP53) and the autism associated protein p21 (CDKN1A) suggesting that the deletion disrupts regulation of the cell cycle and directly impacts expression. Our analyses suggest alterations in the expression of these well-known cell cycle regulators could be mediated by reduced expression of several interacting proteins that map to the deletion including CDC45, a regulator of DNA replication, TRMT2A, which encodes a known cell cycle inhibitor, as well as LZTR1 a known tumor suppressor. Another notable hub observed in stem cells was that encoding the low affinity nerve growth factor receptor and known NOGO Co-receptor P75, which was increased in expression. The minimal network implicated the NOGO receptor (RTN4R) and the mediator of protein degradation through the proteasome UFD1L, both of which are encoded within the deletion.

In neural progenitor cells (Extended Data Fig. 8c), we continued to see evidence for disruption in NOGO signaling through increased expression of both NOGO (RTN4) and the TRKA receptor, which is associated with autism through rare protein-coding variation and is also a known interactor with P75 and whose signaling is modulated by NOGO signaling. These findings suggest that reduced expression deletion proteins such as the NOGO receptor and less appreciated interacting proteins encoded within the deletion such as PIK4A and ARCV4 are disrupting signaling. Another significant signal emerging from the minimal network in NPCs was for a disruption in RNA metabolism. This was exemplified by a hub centered around The TFIID transcription factor, TAF1 which interacted with the tumor suppressor proteins LZTR1 and LZTS2, the transcriptional activator NFKBIA, an RNA helicase associated with ASD, MOV10, and GNB1L, encoded within the deletion, with roles in cell cycle progression and gene regulation. TAF1 was also connected to the protein-ubiquitination pathways via interactions with HSPA1B and its interactors, both from within and outside the deletion region.

In neurons (Fig. 4g), we identified three major hubs consisting of 1) interactors of the activity-dependent transcription factor *JUN*, including the proteasome subunit PSMD12 and the kinesin KIF2A, both associated with NDD, and BANP, a cell cycle regulator, along with several proteins encoded in the 22q11.2 interval: TRMT2A, RANBP1, GNB1L, MRPL40, SCL25A1, CRKL, with connections to the transcriptional (POLR2A) and chromatin remodeling (HIRA) machineries; 2) components of the protein ubiquitination / metabolism pathway, including SMAD2, COPS5 and WWP2 along with UFD1L, KLHL22, both encoded within the deletion region; and 3) synaptic vesicle trafficking, including CLTCL1 encoding clathrin, the synaptobrevin VAMP2, which is associated with NDD, and SNAP29 located in the 22q11.2 locus and encoding a synaptosome associated protein (Fig. 4g). Overall, our analyses support the notion that multiple distinct but connected pathways are at the core of the transcriptional changes that we observe in deletion carrier neurons: activity-dependent gene expression, protein homeostasis, and synaptic biology.

### Enrichment of synaptic and protein homeostasis ontologies in deletion altered transcripts

We next wondered how changes in gene expression caused by the 22q11.2 deletion might impact neurobiological processes. To this end, we employed recently reported synaptic gene ontologies^31^ to search for potentially converging synaptic biology among the genes differentially expressed in 22q11.2 patient neurons. Strikingly, 239 of the 2,864 transcripts with increased abundance in 22q11.2 neurons possessed a synaptic process annotation in SynGO^31^ (p=1.1 × 10^-10^), with a particular enrichment for transcripts with presynaptic functions in synaptic vesicle cycle (GO:0099504, p_FDR adj_=6.12 × 10^-9^, Fig. 4a, Table S9), while 35 of the 1,328 downregulated transcripts, including five cis genes, had a SynGO annotation. We next wondered whether these 239 synaptic genes were a major contributor to the schizophrenia heritability enrichment we detected in the overall set of transcripts induced in deletion neurons. Indeed, we found a marked reduction in the per SNP heritability for schizophrenia after removing these 239 transcripts from the 2,864 that showed increased abundance in 22q11.2 deletion neurons (Fig. 4b). This reduction was greater than that observed when randomly drawn lists of 239 transcripts were removed from the overall pool of 2,864 more abundant transcripts, suggesting that this modest number of synaptic transcripts explained proportionally more of the heritability than the rest.

A further gene ontology enrichment analysis revealed that genes induced in 22q11.2 neurons were significantly enriched for functions particularly in the protein ubiquitination pathway (GO:0000209, 87 genes, OR=2.13, q = 7.5 × 10^-8^) and with the largest individual enrichment for regulation of synaptic vesicle exocytosis (GO:2000300, 14 genes OR=4.0, q=2.3 × 10^-4^) (Table S10). This enrichment with functions in protein homeostasis and synaptic signaling was specific for induced genes in neurons. Conversely, in the genes induced in earlier developmental stages, the enriched functions were related to developmental processes, including tube morphogenesis and development, along with cell motility, migration and differentiation in hPSCs and embryonic development and cardiac epithelial to mesenchymal transition in NPCs. In comparison, genes reduced by the deletion in neurons highlighted exclusively functions in cilium assembly (GO:0060271, 54 genes, FC= 2.2, q = 6.4 × 10^-5^), while genes reduced in hPSCs and NPCs were not enriched for any biological processes (Tables S11-S13). Together the results of our gene ontology and PPI analyses converge on the same key pathways that are regulated by the 22q11.2 deletion in each cell type. These results further demonstrate that the cell type-specific effects of the deletion involve distinct biological functions that may have clinical relevance for the phenotypic presentation in patients.

### Enrichment for programs associated with activity dependent gene expression

To further investigate which cellular programs might mediate the changes in synaptic gene expression and protein homeostasis observed upon 22q11.2 deletion, we carried out motif enrichment analysis on the genes upregulated (p <0.05) in deletion carrier neurons to identify transcription factor binding motifs that are enriched in this gene set. The motif that was most significantly enriched was for binding of the *JUN*/*FOS* transcription factors (1.6-fold enrichment, p = 10^-14^; Fig. 4c, Table S14). The *JUN* and *FOS* transcription factors are immediate early genes that are activated in response to neurotransmitter release and activate a downstream “activity-dependent” transcriptional cascade to regulate downstream programs, such as protein homeostasis and synaptic transmission^48^.

Notably, there was significant overlap (p = 5.57 x 10^-16^) between the genes altered in deletion neurons that had synaptic ontologies (Table S9) and the altered genes that were targets of *JUN* /*FOS* (Table S14) suggesting that activity dependent gene expression downstream of *JUN/FOS* is a contributor to the synaptic signal that we detected in 22q11.2 deletion neurons. Additionally, a further gene ontology enrichment analysis of the unique *JUN*/*FOS* targets we identified (Table S14) revealed an enrichment of components of the protein ubiquitination pathway (GO:0016567, 29 genes, OR=2.9, p_FDR adj_= 0.00098, Table S15).

Furthermore, transcript levels of *MEF2C*, an activity-dependent transcription factor acting upstream of the JUN / FOS signaling pathway to regulate the expression of immediate early genes^48^, are increased in 22q11.2 deletion carrier NPCs in our discovery dataset (Table S2, Fig. 4d, and validated by qPCR and immunoblotting, Extended Data Fig. 2f,g). *MEF2C* has been shown to negatively regulate synaptic transmission by restricting the number of excitatory synapses^66,67^ . Additionally, *TBX1*, a transcription factor located in the 22q11.2 deletion region and significantly downregulated in these same NPCs (Extended Data Fig. 2d,e), is a known repressor of *MEF2C*^49,50^. Thus, decreased *TBX1* levels due to loss of a copy of 22q11.2 likely result in de-repression of the *MEF2C* transcription factor, a regulator of the *JUN/FOS* signaling pathway, which in turn might reduce synaptic transmission.

Taken together, these results indicate that activity dependent gene expression is changed in deletion carrier cells, likely impacting downstream protein homeostasis and synaptic transmission.

### Reduced network activity in 22q11.2 deletion neurons

Overall, our data suggests that changes linked to the 22q11.2 deletion during the development of excitatory neurons alter the balance of the *JUN/FOS* transcriptional pathway, which has well established roles in activity dependent gene expression^48^. We thus hypothesized that the transcriptional activation of this pathway and its targets, which plays a role in reducing synaptic transmission upon sustained activity^48^ might result in decreased network activity in neuronal cultures with 22q11.2 deletion.

We thus asked whether neurons from 22q11.2 deletion carriers exhibited changes in network activity. Previously, we had shown that by 42 days of excitatory differentiation, neurons derived from control cell lines were spontaneously active and that their rate of firing was governed almost entirely by network activity mediated through synaptic connectivity^34^. We used multielectrode arrays (MEAs) to monitor neuronal network development and activity over 42 days of neuronal differentiation^34^. In neurons derived from patients with 22q11.2 deletion, we detected a significantly lower spiking rate from 21 days of differentiation and onward, when compared to controls (N = a total of 162 wells from 21 cell lines) (Fig. 4e,f). We found this result striking, as it was consistent with the notion that the altered abundance of synaptic transcripts and activity-dependent gene expression we observed by RNA sequencing was associated with functional effects on network activity in 22q11.2 deletion neurons.

### Gene editing of the 22q11.2 deletion

To complement our patient driven study and assess whether the 22q11.2 deletion was sufficient to explain the transcriptional changes we observed in our patient-based discovery cohort, we used CRISPR/Cas9 to engineer the 22q11.2 deletion in a human embryonic stem cell line (H1/WA01). Using guide RNAs that cut within the low copy repeats (LCRs) flanking the 3Mb 22q11.2 deletion, we generated heterozygous 22q11.2 deletion cell lines at a very modest frequency (2/1000), as well as many non-targeted but otherwise isogenic controls (Fig. 5a-d). We then subjected the two deletion clones and two non-targeted control clones to neuronal differentiation and performed RNA sequencing at the same differentiation stages we assessed previously (d0 hPSCs, d4 NPCs and d28 excitatory neurons). In PCA, components one and two separated each of the samples by differentiation state, with the stem cell, NPC and neuronal cell lines showing strong reproducibility of differentiation across replicates (Fig. 5e). Impressively, components three and four then separated each of the samples based on their deletion status, with 22q11.2 deletion samples substantially separated from their non-targeted counterparts (Fig. 5f, Extended Data Fig. 9a). This separation was not solely due to deleted cis genes as it persisted upon removal of these genes from the PCA, indicating that it was a more global phenomenon in the transcriptome of the edited lines. Importantly, the genes driving the separation in PC3 and PC4 were largely shared by those detected differentially expressed in the discovery cohort. Out of the top 100 negative and positive loadings for PC3, 79 and 83, respectively, were nominally significantly changed also in neurons in the discovery cohort (p < 0.05). For PC4, this overlap was 39 and 60 out of 100, for negative and positive loadings, respectively.

**Fig. 5.**
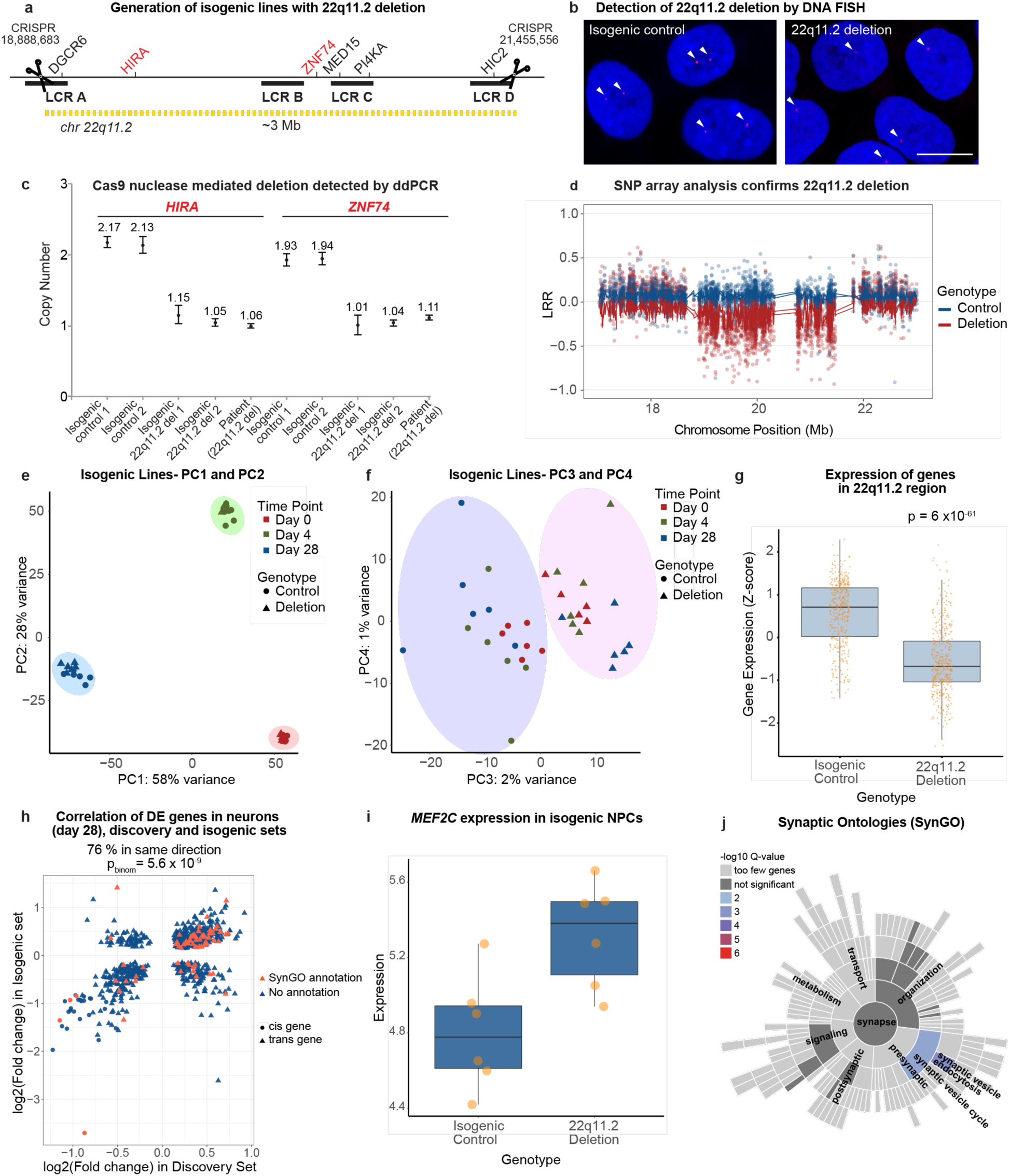
Validation of causality between the differentially expressed genes and the deletion genotype in an isogenic setting. **a**, Generation of isogenic lines with 22q11.2 deletion using CRISPR Cas9 guide RNAs that cut within the low copy repeats (LCRs) flanking the 3Mb 22q11.2 deletion. The coordinates for the genomic position of the CRISPR guides on chromosome 22 are indicated (Hg19). **b**, Detection of isogenic 22q11.2 deletion using DNA FISH analysis and a probe generated probe using CTD-2300P14 (Thermo Fisher Scientific, Supplier Item: 96012). Blue = DAPI (DNA), Red=22q11.2 region. Scale bar: 10um. **c**, ddPCR assay to determine the copy numbers of the HIRA and ZNF74 genes, located in the 22q11.2 region, to validate isogenic deletion of 22q11.2. **d**, SNP array marker intensity (LRR) for SNPs overlapping the deletion locus confirms isogenic 22q11.2 deletion in two clones (red). **e,f**, Principal component analysis of cell lines with and without isogenic 22q11.2 deletion. Circles = genes within the 22q11.2 interval (cis). Triangles = genes outside 22q11.2 (trans). **e**, PC1 and PC2 separate cells by developmental stage. **f**, PC3 and PC4 separate cells by deletion genotype. **g**, Significant downregulation of genes in 22q11.2 region in lines with isogenic 22q11.2 deletion. **h,** Correlation of fold changes in differentially expressed genes in discovery and isogenic datasets in neurons. 32 genes were detected and significantly changed in transcript levels in the discovery cohort and the isogenic lines (adjusted p-value < 0.05 in both experiments), of which nine were located outside the deletion region (*FAM13B*, *KMT2C*, *HYAL2*, *DNPH1*, *ZMYM2*, *VAPB*, *SMG1*, *CPSF4*, *MAP3K2*) and the rest were in cis. All 32 genes were changed in the same direction both in the discovery and isogenic cohorts (p = 0.004, binomial test). Genes with a SynGO annotation are shown in red, genes with no SynGO annotation are shown in blue. Circles = cis genes. Triangles = trans genes. **i**, *MEF2C* is upregulated in NPCs of deletion carriers compared to isogenic controls, similar to the discovery dataset. **j**, SynGo annotation of genes induced in isogenic neurons with 22q11.2 deletion showing enrichment for synaptic vesicle cycle and endocytosis.

We next proceeded to perform differential expression analysis to delineate transcriptional changes present in clones edited to contain the 22q11.2 deletion (Tables S16-S18). As expected, the edited lines showed systematic downregulation of genes in the deletion region at all cell stages (p= 6 x 10^-61^, Mann-Whitney test) (Fig. 5g) with 26, 25, and 29 deleted genes passing individually FDR< 5% cutoff in the isogenic hPSCs, NPCs, and neurons. This further confirmed successful introduction of the heterozygous 3Mb deletion in this background. Notably among these and like the discovery set, *CAB39L* was consistently upregulated at all differentiation stages in lines with isogenic 22q11.2 deletion. Overall, we also observed a highly significant number of genes exhibited aligned changes in transcript abundance between the discovery cohort and edited samples (p<0.05) across all differentiated stages analyzed: hPSCs, 75% (200 out of 268 p=3 x 10^-16^, binomial test); NPCs, 83% (124 out of 150 p=1.7 x 10^-16^, binomial test) and neurons, 76% (604 out of 791p=5.6 x 10^-9^, binomial test) with strongly correlated effect sizes (r_hPSC_=0.7, r_NPC_=0.82, r_neuron_= 0.56, Pearson correlation); (Fig. 5h, Extended Data Fig. 9b,c).

We next wondered whether the pathways and cellular programs that were altered in a cell-type specific manner in our discovery dataset were also altered in the edited lines. To this end, we examined the expression of genes contributing to the minimal PPI networks identified at each cell stage in the discovery dataset (Fig 4g and Extended Data Fig. 8) and found that an overwhelming majority of these genes are changed in the same direction in cells with isogenic 22q11.2 deletion at each stage, with 90%, 88% and 86% of the genes contributing to the PPI network in stem cells, NPCs and neurons respectively, being altered in the same direction in the isogenic dataset compared to the discovery dataset. Notably, the activity dependent gene *MEF2C* was also increased in NPCs of H1 deletion carrier cells compared to isogenic controls (Fig. 5i, Extended Data Fig. 9d,e).

Furthermore, upon synaptic process annotation in SynGO we observed a replication of the induction of genes (p < 0.05) involved in synaptic vesicle cycle and endocytosis in the edited neurons with 22q11.2 deletion (GO: 0099504, p_FDR adj_ = 0.0029) (Fig. 5j, Table S19). Overall, of the 239 transcripts with synaptic functions in the discovery dataset (Fig. 4a), 49 were also more abundant in neurons (p < 0.05) harboring the engineered 22q11.2 deletion (Expected = 39 genes, p<0.012, binomial test), out of which 21 passed the FDR < 5% cutoff for significance.

Additionally, the 87 transcripts implicated in the ubiquitination pathway that we found to be more abundant in 22q11.2 deletion carrier neurons were on average 0.29 standard deviations (SDs) higher expressed in the edited lines (95%-CI:0.18-0.41 SDs, p =3.9×10^-7^, t-test). Moreover, 19 of these transcripts were individually significantly (FDR < 5%) more abundant after gene editing of the deletion (p=0.03 binomial test) supporting a causative connection between the deletion genotype and altered transcript abundance for components in the ubiquitin-proteasome system in neurons. Furthermore, 28 out of the 99 JUN target genes induced in the discovery dataset were also induced in neurons with isogenic 22q11.2 deletion (p<0.05) (p = 0.00046, binomial, expected overlap = 14 genes). Finally, encouraged by the replication of the differential expression signal in the edited deletion lines, we examined these genes (p<0.05) for association to schizophrenia. Remarkably, variants surrounding the induced genes in the edited lines revealed significant gene-wise association to schizophrenia consistent with the observation in the discovery cohort (β=0.11, SE=0.029, p = 6.6×10^-5^, N=1611 genes, MAGMA). Thus, we conclude that the 22q11.2 deletion is indeed sufficient to explain most transcriptional effects we found to be associated with the deletion in our case-control cohort, including those related to the genetic risk for schizophrenia.

### Reduced pre-synaptic protein abundance in 22q11.2 deletion neurons

As an independent means of examining whether the 22q11.2 deletion impinged on presynaptic components in excitatory neurons, we performed whole cell proteomics on day 28 neurons from two patients and two controls (Fig. 6a) As expected, peptides mapping to genes within the 22q11.2 interval were reduced in neurons harboring the deletion relative to levels in controls (Fig. 6b; Table S20).

**Fig. 6.**
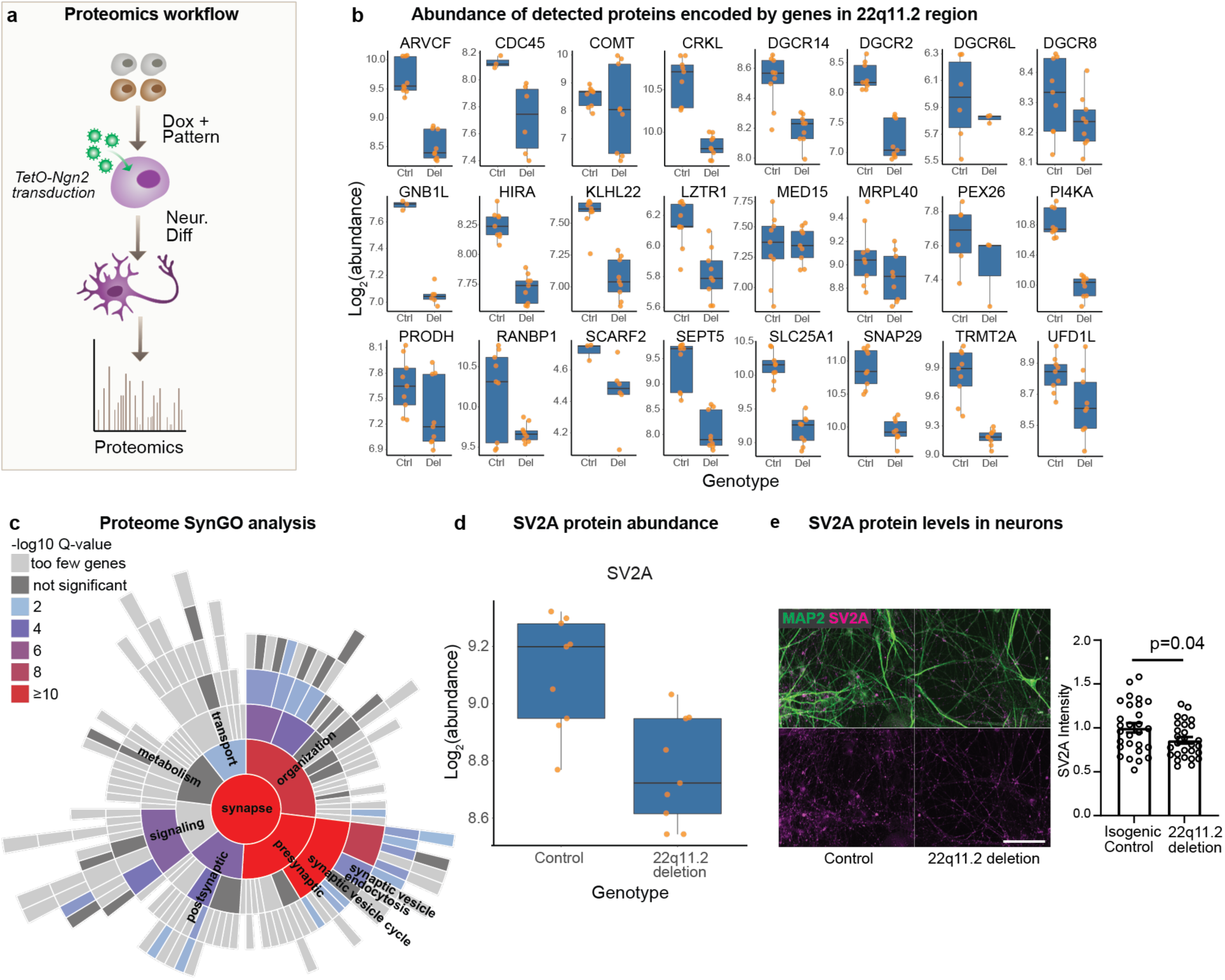
Whole cell proteomics on 22q11.2 deletion neurons. **a,** Workflow schematic. Neurons from deletion carriers and controls were harvested 28 days post neuronal induction. **b**, Abundance of proteins encoded by genes in the 22q11.2 region detected by proteomics in neurons. Del = 22q11.2 deletion. Ctrl = control. **c**, Synaptic gene ontologies (SynGO) in proteins downregulated in deletion carrier neurons. **d**, SV2A protein levels detected by proteomics are decreased in deletion carrier neurons. **e**, SV2A protein levels detected by antibody staining are decreased in day 28 neurons derived from isogenic lines with 22q11.2 heterozygous deletion compared to controls. (Left) Representative confocal images of control and 22q11.2 deletion neurons stained with antibodies against SV2A (magenta) and MAP2 (green). Scale bar is 100 µm. (Right) Quantification of total SV2A fluorescence within MAP2-positive neurites normalized to isogenic controls. Data are means ± SEM. Individual points are analyzed fields of view from 4 independent inductions per condition. Statistical analysis by Student’s t test reveals statistically significant (p=0.037) decrease in SV2A levels in deletion neurons.

Importantly, consistent with the altered expression of activity-dependent genes (Fig. 4c,d,g and Table 14), and the reduced synaptically-driven network activity in 22q11.2 deletion neurons (Fig. 4 e,f), we found that proteins downregulated in 22q11.2 deletion neurons were enriched for synaptic gene ontologies (Fig. 6c). In total, 184 of the proteins that were downregulated in deletion carrier neurons had SynGO annotations. Of these, 68 were upregulated at the transcriptional level. Additionally, 31 proteins were upregulated in deletion carrier neurons and had SynGO annotations; 4 of which were also upregulated at the mRNA level (Extended Data Fig. 9f).

The synaptic components exhibiting alterations in deletion neurons were predominantly presynaptic and specifically involved in synaptic vesicle cycle (p_FDR adj_ = 3.5×10^-19^) (Fig. 6c; Table S21), and included Synaptotagmin 11 (SYT11), Neurexin-1 (NRXN-1), and Synaptic Vesicle Glycoprotein 2A (SV2A). SV2A (Fig. 6d) regulates vesicle exocytosis into synapses and works in presynaptic nerve terminals together with Synaptophysin and Synaptobrevin^68,69^. We note this finding also converges with genetic studies as rare variants in SV2A have been shown to be significantly associated with schizophrenia^65,70^. Similarly, *NRNX1* has established roles in schizophrenia^65,71,72^ and *SYT11*, located on the chromosome locus 1q21-q22 may be a risk gene for schizophrenia^73^. We confirmed the decreased expression of SV2A (Fig. 6e), along with the reduction of protein levels of SYT11 (Extended Data Fig. 9g) and NRXN1 (Extended Data Fig. 9h) in 22q11.2 deletion neurons by immunostaining or immunoblotting. Additional proteins with schizophrenia rare variant associations (via the SCHEMA consortium^65^) altered in 22q11.2 deletion neurons included DNM3, MAGI2 and TRIO (downregulated in patient neurons) and HIST1H1E, SRRM2 and ZMYM2 (upregulated in patient neurons) (Table S21).

## Discussion

Here we have explored the transcriptional and functional consequences of the 22q11.2 deletion on human neuronal differentiation. Our findings lead to several new insights into the biology of 22q11.2 deletion syndrome and how it confers risk for the development of varied psychiatric disorders as neural development and differentiation unfold. Notably, we found that the genes whose expression is perturbed in deletion carriers directly connect the effects of the deletion on neuropsychiatric phenotypes to genes and pathways implicated in NDD, ASD and schizophrenia through prior large-scale exome sequencing and GWAS studies^27,28,32,45–47,63,64,74^. Thus, rather than working through independent mechanisms, our studies suggest the deletion confers risk for these various conditions at least in part by converging on the same gene products and pathways that are more widely disturbed in other patients.

We used a new tool that we developed and report here to ask which minimal PPI networks best explain the changes in gene expression we observed. This analysis revealed that a surprising number of deletion components likely play a role in the transcriptional signals. We therefore propose a model in which reduced abundance of multiple factors within the deletion region leads to highly distributed effects on many genes outside the deletion. Through the course of development, the deletion affects distinct sets of genes. In stem cells and neuronal progenitor cells the deletion impacts pathways linked to proliferation, NOGO signaling and RNA metabolism. In neurons, the deletion alters activity-dependent gene expression, protein homeostasis and ultimately, presynaptic biology. Overall, it was notable that *MEF2C*, an activity dependent transcription factor and negative regulator of excitatory synaptic density^66,67^ is overexpressed in NPCs with 22q11.2 deletion, likely due to the loss of one copy of *TBX1*, a known *MEF2C* inhibitor located in the 22q11.2 interval^49,50^. Increased expression of *MEF2C*, could, in turn, lead to premature activation of the JUN and FOS pathway, which would be predicted to result in reduced network activity and synaptic connectivity.

To directly test this idea, we examined whether neurons from 22q11.2 deletion carriers displayed reduced synaptic functionality. Using a network activity assay in these cells, which we have previously shown was largely driven by a mixture of AMPA and NMDA receptor mediated transmission^34^, we indeed found this to be the case. Many of the patients’ neurons showed a significant overall reduction in network activity relative to controls. Thus, the deletion was not only associated with induction of activity dependent gene expression, but also associated with aligned changes in neuronal function. Based on these findings, we would thus expect a decreased expression of synaptic proteins, which we do, indeed, detect.

Our proteomic examination of 22q11.2 deletion neurons afforded an orthogonal examination of synaptic components in these cells and independently identified significant presynaptic alterations, including alterations in components that we could not ascertain by RNA sequencing such as the schizophrenia associated gene SV2A, a key mediator of pre-synaptic function. This last result is of translational and therapeutic importance given the existence of a positron emission tomography (PET) radiotracer specific for SV2A based on the drug Levetiracetam which now enables the *in vivo* investigation of presynaptic protein levels in the patient brain^75^. Interestingly, a recent PET-imaging study utilizing this SV2A radiotracer found a significant reduction in the abundance of this presynaptic component in the cortex of schizophrenia patients relative to controls^76^. Careful genotyping of this schizophrenia patient population was not carried out prior to imaging and our results suggest that a more specific study examining SV2A levels in 22q11.2 deletion carriers of varying diagnoses would be warranted.

Early during neuronal differentiation, we found that a significant number of the genes differentially expressed in deletion carriers had been previously linked to damaging or LoF sequence variants more widely identified in NDD and ASD. This enrichment for overlap between broader genetic signals in ASD and the effects of the 22q11.2 deletion was very significant when we considered the known biochemical interaction partners of gene products implicated in ASD. These findings are consistent with smaller scale studies investigating transcriptional effects of individual genes, such as *FOXP1* or *CHD8*, linked with autism, and found to regulate the expression of ASD-relevant pathways ^77,78^.

Interestingly, as differentiation proceeded and cells took on a post-mitotic, excitatory neuronal identity, the effects of the 22q11.2 CNV on expression of genes outside of the deletion lost enrichment for genes implicated in NDD/ASD and acquired an enrichment for genes harboring rare inactivating exome variants preferentially associated with schizophrenia. The influence of the 22q11.2 deletion on expression of neuronal genes associated with schizophrenia was not limited to those impacted by rare schizophrenia mutations acting with large effect. We also found that the deletion affected neuronal genes that were in linkage disequilibrium with common genetic variants associated with schizophrenia, a result replicated using genotypic data from two independent GWAS studies. Just as signal from ASD/NDD associated genes was absent in the neuronal stage of differentiation, the enrichment for effects on schizophrenia associated genes was absent in stem cells and NPCs. This surprisingly selective signal is likely to reflect stage-specific cellular programs, such as synaptic processes (for example those listed in Tables S9, S19 and S20) being specific to neurons.

We found these transcriptional results striking as NDD and ASD are linked to biological processes acting early in brain development^79^, while sequence variants associated with schizophrenia have been previously shown to be enriched for genes expressed in excitatory neurons and more recently for genes functioning in excitatory synaptic transmission^80^. It is important to note that our findings were not merely the result of looking at a chance list of genes in cell types clearly impacted in these diseases. While we did find that the overall gene expression profile of our excitatory neurons was enriched for expression of genes implicated in schizophrenia, the specific transcripts induced by the 22q11.2 deletion showed significantly greater enrichment in all tests we performed. Thus, we hypothesize that by looking in a human cell type with disease relevant biology, we were able to identify previously unappreciated effects of the 22q11.2 deletion.

Overall, our findings support human genetic studies suggesting that neuropsychiatric CNVs such as 22q11.2 deletion likely interact with risk variants in the genetic background^23–25^. Transgenic mice carrying syntenic deletions that model the human 22q11.2 deletion have produced a wealth of datasets around neurodevelopmental abnormalities linked to the deletion or to individual genes within the region^13,20,81^. It is, however, important to keep in mind that such transgenic mice do not have genetic backgrounds harboring human polygenic risk alleles, which explain the majority of heritable variation in schizophrenia and other psychiatric phenotypes^47^. Therefore, while non-human model systems offer invaluable biological insight, they fall short of reproducing human specific gene regulatory effects underlying complex human disorders.

Individual genes within the 22q11.2 region have been at the center of several studies aiming to identify causal genes underlying the 22q11.2 deletion syndrome. Several of these studies, using rodent, and more recently, human^41^ models, have reported defects in synaptic processes and brain connectivity ^82–84^, many with a focus on *Dgcr8*, which encodes a subunit of the microprocessor complex which mediates microRNA biogenesis^13^. Khan et al^41^ identified a calcium signaling defect in organoids containing mixed cell types derived from 22q11.2 deletion and controls individuals, which could then be rescued by *DGCR8* overexpression. Whether these phenotypes can be recapitulated with a scaled sample set and defined cell types remains to be seen. At face value, alterations in *DGCR8* might seem like a promising candidate for the distributed effects on gene expression we observed across many transcripts. However, reduced microRNA function from lower *DGCR*8 copy number would predict an increased rather than decreased abundance of the synaptic proteins we found. Another candidate, *DGCR5*, which encodes a long non-coding RNA within the 22q11.2 interval, has previously been shown to regulate several transcripts encoding genes associated with schizophrenia^85^. However, that study found that reducing the function of *DGCR5* lead to a reduction in the expression of its targets, again the inverse of our finding.

A challenge in studying psychiatric conditions has been that it is difficult to establish causal relationships between genetic variants of interest and their effects. In this study we utilized CRISPR/Cas9 to generate the 22q11.2 deletion in a control human stem cell line by inducing double strand breaks within the same repetitive elements that are normally important mediators of the deletion. While the process was relatively inefficient, we were able to obtain two independent clones that carried this large structural variant on one of the two alleles. Using these edited cells, we could then ask, without confounding by inherited variation elsewhere in the genome, which associations we had previously observed was the deletion sufficient to cause. We found that the deletion in this isogenic setting was sufficient to induce significant and aligned alterations in the expression of genes contributing to the minimal PPI network at each of the three differentiation stages, including changes, in neurons, in the activity-dependent, presynaptic, and ubiquitin/proteasome pathways as well as the heritability enrichment for schizophrenia.

When combined with genetic findings from 22q11.2 patients^23–25^, our observations lead us to a model in which the 22q11.2 deletion exerts a strong effect on genetic risk factors for NDD and ASD genes early in differentiation, while in more differentiated neurons the gene regulatory influence of the deletion shifts to risk factors for schizophrenia. Our gene editing experiments suggest that these distinct “pushes” on NDD/ASD and schizophrenia risk occur regardless of one’s genotype.

How exactly the 22q11.2 deletion might regulate the expression of genes outside of the deletion region remains a matter of great interest. Many studies have highlighted the role of miRNAs as possible mediators of some of the phenotypes, particularly given that *DGCR8* is located within the region. However, as discussed earlier, reduced levels of DCGR8 would not explain our finding of reduced synaptic proteins. One intriguing possibility is that 22q11.2 deletion might impact chromatin architecture, thereby regulating the expression of genes outside of the deletion region. Spatial organization of the genome has been shown to play a critical role in cell type-specific regulation of transcription^86^, and structural variants, such as CNVs, have been shown to alter chromatin architecture, leading to disease^87^. The 22q11.2 deletion lacks a large portion of chromosome 22^43^, which might impact chromatin organization. Indeed, a recent study using lymphoblastoid cell lines with 22q11.2 deletion revealed changes in their genome architecture^88^. It is thus possible that the 22q11.2 deletion spatially rearranges the genome of neuronal cells, resulting in mis-regulation of genes linked to neuropsychiatric disorders.

The current study is not without its limitations. Even though it is, to our knowledge, one of the largest of the effect of 22q11.2 deletion on human neuronal cells, our current sample size still falls shorts of enabling us to stratify the cohort by diagnosis, age, sex, or deletion size. Future studies with even larger sample sets could be sufficiently powered to enable the comparison of cells from 22q11.2 deletion patients with or without schizophrenia, or with or without intellectual disability or ASD, for example, to more comprehensively delineate the cellular and transcriptional changes associated with each diagnosis. It would also be interesting to stratify the cohort with respect to deletion size: while the 3Mb deletion is by far the most common, accounting for around 90% of cases, smaller nested deletions within the region still result in similar symptoms and diagnoses^3,7,38,39^. While the current study only includes two such shorter deletions, larger studies could be better poised to identify common and distinct signatures of the distinct deletions. Other interesting co-variates to examine include donor age and sex, which do not appear to drive any of the transcriptional differences and signatures we report here but might result in subtle differences that could be detected with a larger sample set.

Collectively, the novel iPSC lines, CRISPR edited cell lines, RNA sequencing data and functional phenotypes we report here will provide a framework for evaluating future therapeutic targets and candidates for 22q11.2 carriers. These 22q11.2 carriers represent an interesting population for drug discovery as they are a group of individuals with more homogenous, yet still textured risk of these psychiatric illnesses. For instance, with the tools we report here, it should be possible to quantitatively address which combinations of the immediate consequences of the deletion most contribute to various components of the gene expression effects we have observed, including deficits in expression of presynaptic proteins. While these efforts are beyond the scope of our current study, we suggest that as aspects of the gene expression signal we observed are rescued, the functional relevance of such findings could be tested in the context of whether neuronal network activity is also restored in patient neurons. Through this approach, the likely multifaceted contributors to psychiatric illness that the 22q11.2 deletion confers could be quantitatively deciphered and the best approaches for alleviating its effects identified.

## Acknowledgements

We thank the many donors, institutions and investigators world-wide that provided their cell lines and supported the publication of the results. We are indebted to Maura Charlton, Genevieve Saphier and Kristen Elwell for their assistance with the regulatory and logistical efforts required to acquire and sequence hiPSC lines. We regret the omission of any relevant references or discussion due to space limitations. The Genomics Platform at the Broad Institute performed sample preparation, sequencing, and data storage. This work was funded predominantly by U01MH105669 (NIH/NIMH), with additional support from the Stanley Center for Psychiatric Research at the Broad Institute, R37NS083524 and U01MH115727. RN was also supported by a NARSAD young investigator award (Brain and Behavior Research Foundation) and a Bn10 grant (Broad Institute), and OP was also supported by the Sigrid Juselius Foundation, Orion Research Foundation, Instrumentarium Science Foundation, and Päivikki and Sakari Sohlberg Foundation.

## Data Availability Statement

The raw sequence datasets generated during the current study are not currently publicly available due to patient confidentiality and multiple different consents of population cohorts used but subsets of the data are available from the corresponding authors on reasonable request. Computer code relevant to the PPI analysis has been deposited in GitHub (https://github.com/alexloboda/PPItools). Other computer code and data analysis will be made available upon request.

## Author contributions

R.N., O.P. and K.E. conceived the work, designed the experiments, analyzed the data and wrote the manuscript. R.N. supervised and performed the experiments, with help from A.T., C.B., M.T., R.M., E.J.G., V.V., D.H., E.P., and E.Z. O.P. performed the computational analysis, with help from M.T. and G.G. M.A. performed the PPI analysis, with help from A.L. and supervision from M.D. C.B. performed the proteomics experiments with help from J.A.P. and supervision from J.W.H. A.G. carried out the SNP heritability analysis, with oversight from B.N. T.S. carried out the rare variant analysis. J.S. performed the MEA analysis. D.M., A.B., A.M.B. and D.Z.H. carried out the CRISPR editing, supervised by L.E.B. A.N. and C.L. assisted with stem cell compliance and data deposition. O.K., E.H., and M.K. provided the NFID cell lines, with oversight from A.P. C.M.H. and A.K.K. contributed the KI cell lines. B.C. and D.M. provided cell lines from Mclean Hospital. J.M., R.A. provided the Umea samples, R.D. provided the Stanford cell lines, and R.P. provided the MGH cell line. S.M. and S.H. provided guidance throughout the project.

---

**Extended Data Fig. 1.**
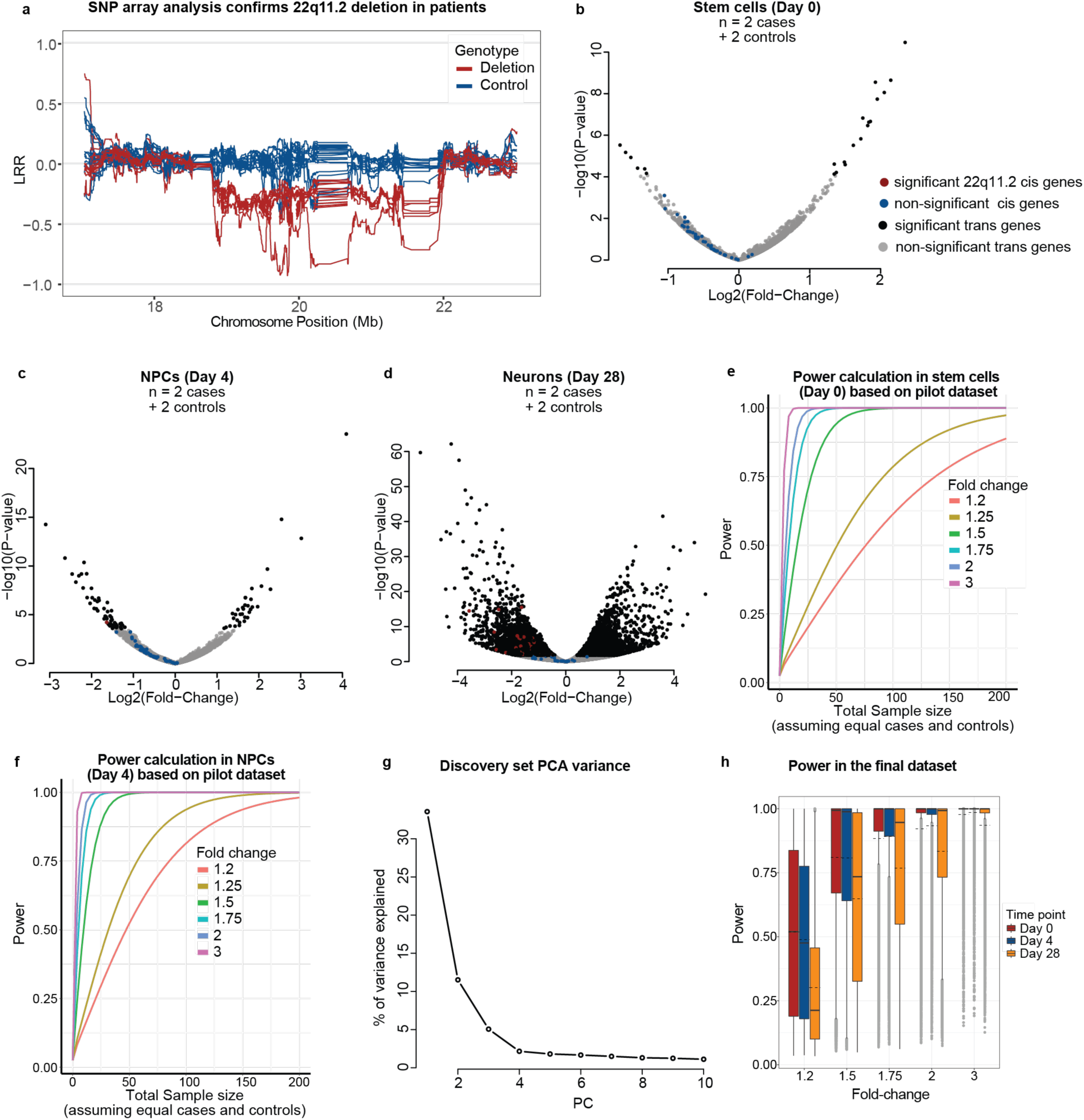
Discovery and pilot datasets. **a**, Validation of the full-size deletion in 22q11.2 lines used in the current study by sliding-window average of SNP marker intensity (LRR) in the deletion locus. **b-d**, Volcano plots showing differentially expressed genes in the pilot dataset. **e**, Power estimation in the pilot dataset for median expressed genes for different fold-changes and sample sizes in stem cells. **f**, Power estimation in the pilot data set for median expressed genes for different fold-changes and sample sizes in neuronal progenitor cells (NPCs). **g**, Variance explained by the first 10 principal components in the discovery sample. **h**, Estimated power in the final (discovery) dataset, at each time point.

**Extended Data Fig. 2.**
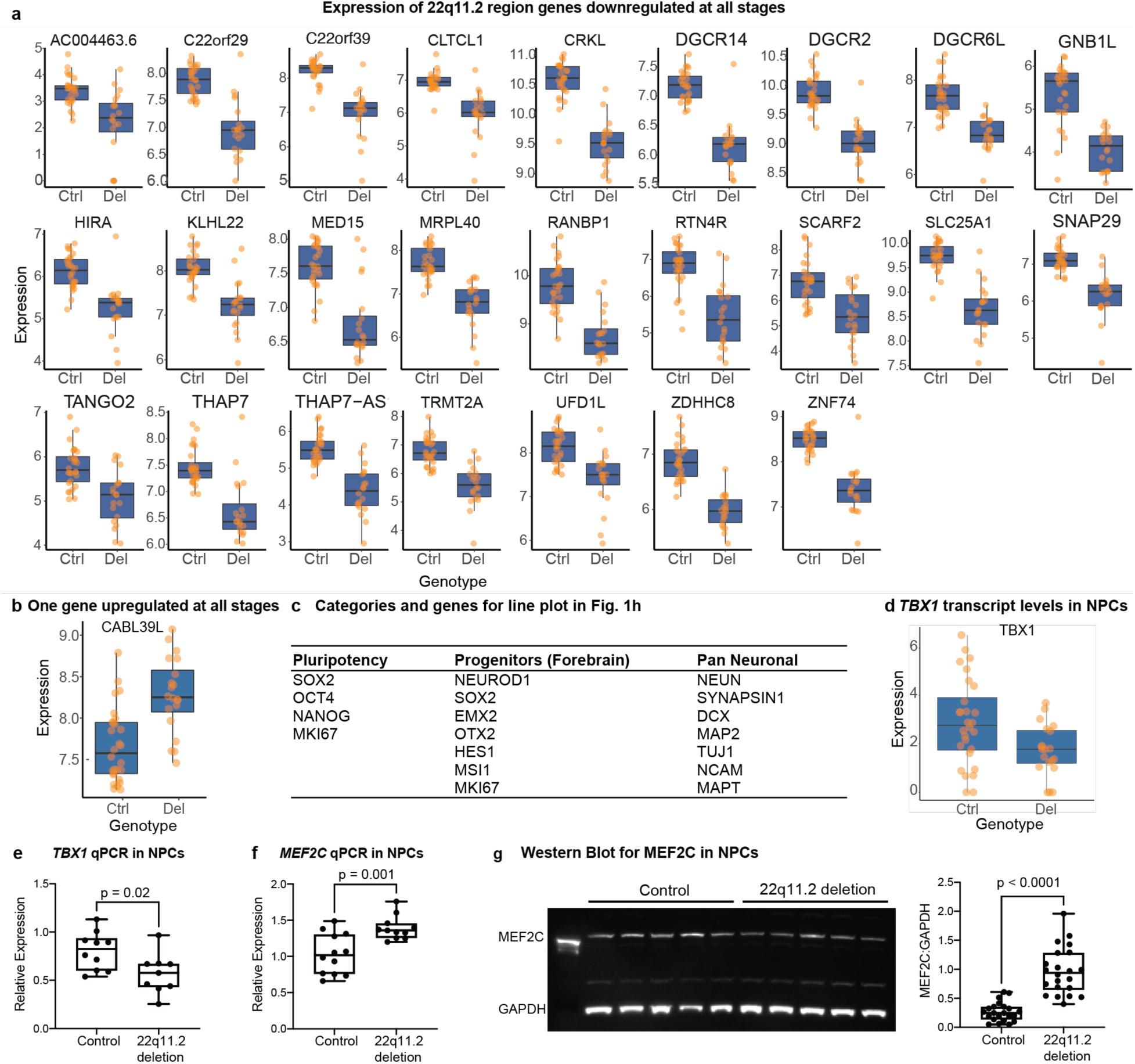
Expression of differentially regulated genes. **a**, Expression of significant cis genes shared across all three developmental stages. **b**, *CAB39L* is the only trans gene upregulated in all developmental stages. **c**, List of categories and genes used in Fig. 2a. **d**, *TBX1* is downregulated in NPC of 22q11.2 deletion carriers. **e**, Relative expression of *TBX1* via qPCR in Day 4 NPCs from control and 22q11.2 deletion patients (Samples: 3/3, 2BR/2TR, p<0.05). **f**, Relative expression of *MEF2C* via qPCR in NPCs from control and patients (Samples:3/3, 2BR/2TR, p<0.01). **g**, Expression of MEF2C in total protein lysates from control and 22q11.2 deletion NPCs. (Left) Total protein lysates from control (left five lanes) and deletion lines (right five lanes) probed for MEF2C (top) and GAPDH (bottom). (Right) Statistical analysis by Student’s t test reveals statistically significant decrease in MEF2C expression in the deletion lines. (Samples: 5/5, 1BR/3TR, p<0.0001). BR = biological replicate (independent differentiations); TR = technical replicate (independent wells).

**Extended Data Fig. 3.**
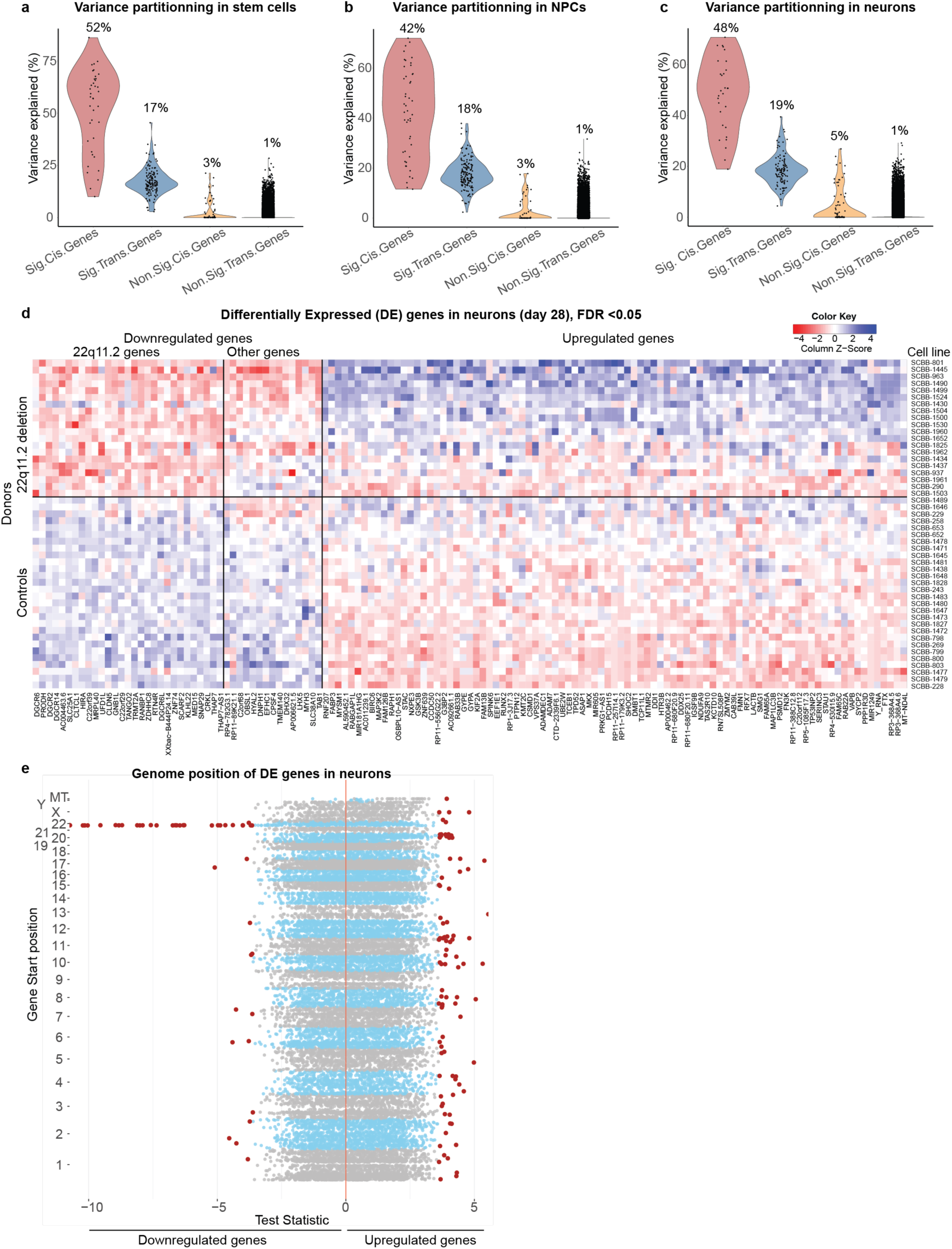
Variance partitioning and expression of differentially regulated genes. **a-c**, Variance in gene expression explained by the deletion genotype in different gene categories in the final dataset in **a**, Stem cells, **b**, Neuronal progenitor cells and **c**, Neurons. **d**, Heatmap of 133 genes differentially expressed in neurons showing the range of expression, in all donor lines, of genes down or upregulated. **e**, Test statistic for differential expression plotted by chromosomal position of differentially expressed genes in cells with 22q11.2 deletion. Differentially expressed genes (FDR<5%) are colored in red.

**Extended Data Fig. 4.**
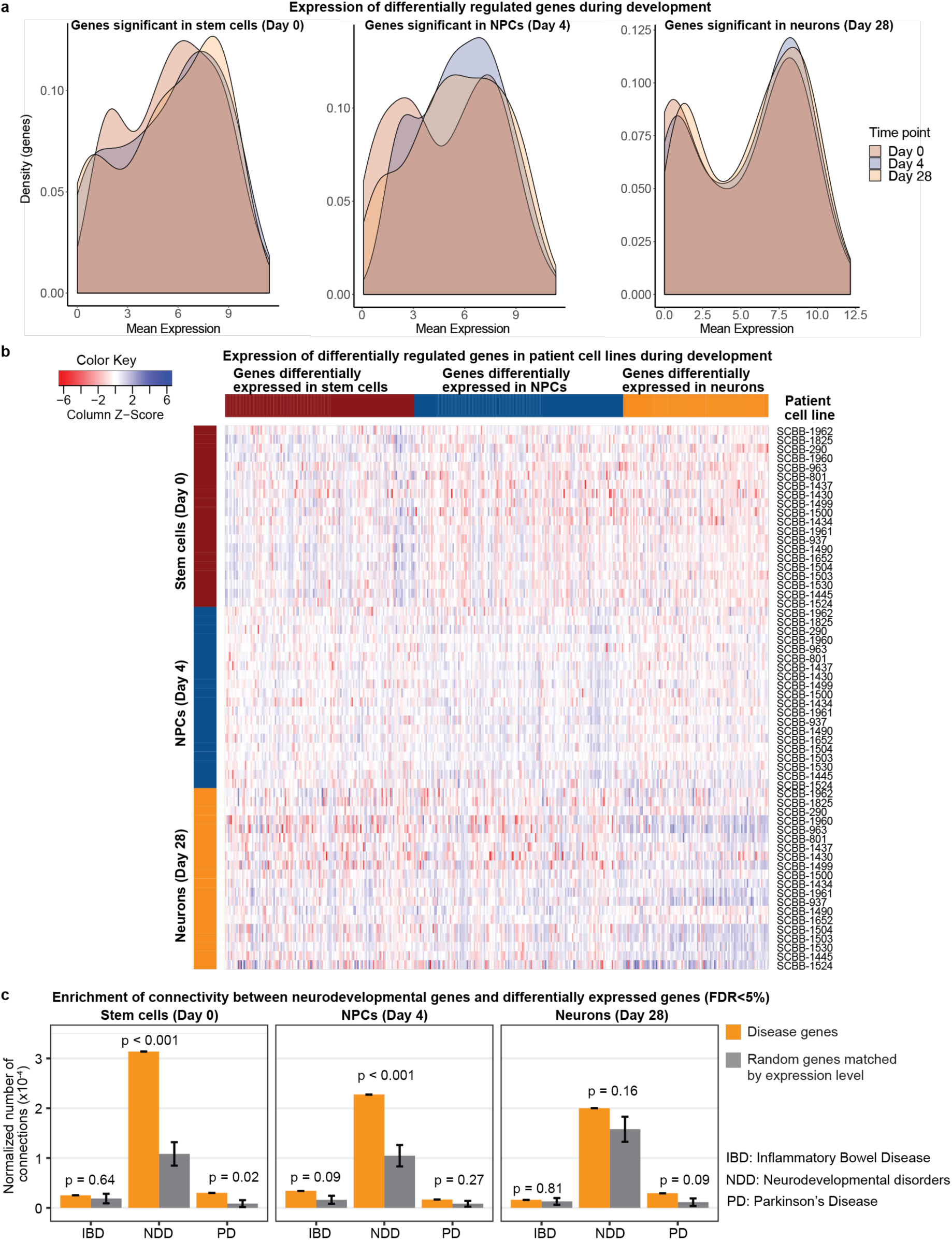
Expression of genes differentially regulated at specific cell stages in 22q11.2 deletion cells across developmental stages. **a**, density plots and **b**, heatmap showing that the differentially regulated genes are expressed at similar levels across all three cell stages. **c**, Connectivity enrichment analysis. Enrichment of interactors of proteins encoded by genes associated with intellectual disability and autism (NDD) and differentially expressed genes (FDR<5%) in the early developmental stages after 1000 random permutations. Gene products encoded by genes linked to inflammatory bowel disease (IBD) yielded no enrichment for protein-protein interactions between the differentially expressed genes at any cell stage, and genes linked to Parkinson’s disease (PD) yielded no enrichment in NPCs or neurons.

**Extended Data Fig. 5.**
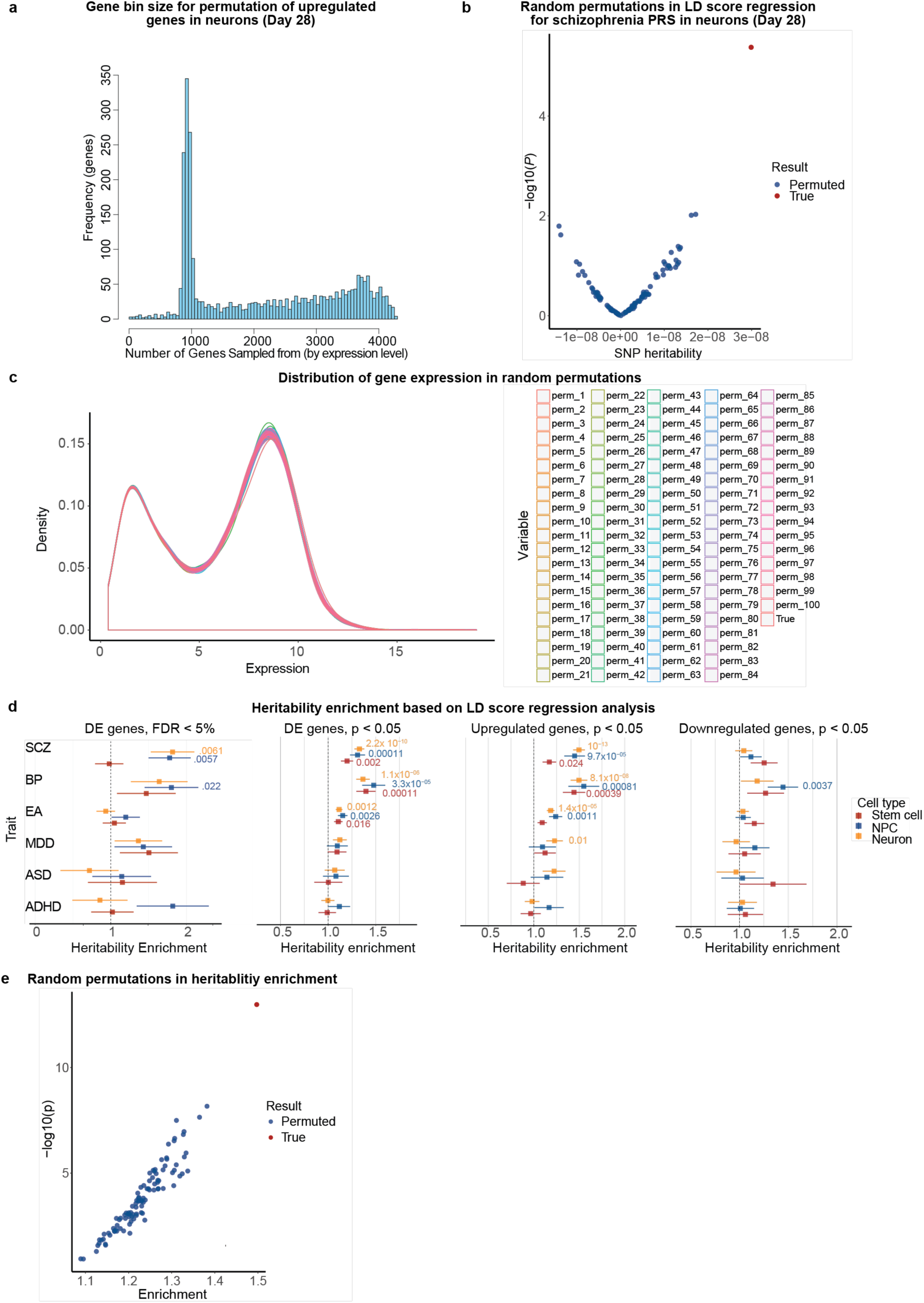
LD-score regression analysis. **a**, Gene bin size for permutation of upregulated genes in neurons. **b**, Per-SNP heritability enrichment in LD score regression for random permutations (in blue) and the upregulated genes (in red) in neurons **c**, Distribution of gene expression in random generated gene sets in the 100 permutations for per-SNP heritability. **d**, Heritability enrichment analysis of six traits across the three developmental cell stages. SCZ= schizophrenia, BP=bipolar disorder, EA= educational attainment, MDD=major depressive disorder, ASD=autism spectrum disorder, ADHD=attention deficit hyperactivity disorder. **e**, LD score regression heritability enrichment in random expression matched gene lists from 100 permutations (in blue) compared to the up-regulated genes in neurons (in red).

**Extended Data Fig. 6.**
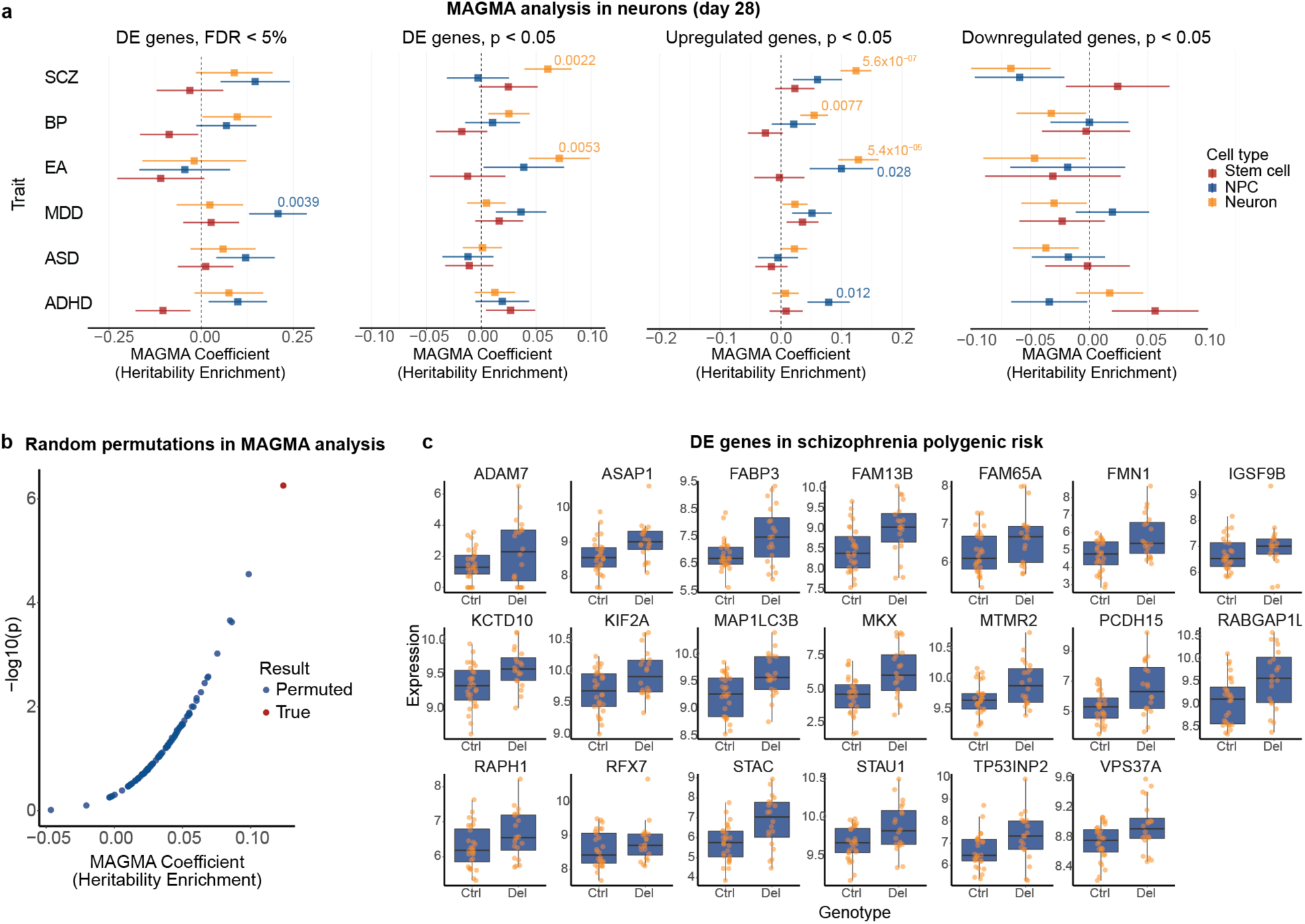
MAGMA (Multi-marker Analysis of GenoMic Annotation) analysis in neurons. **a**, MAGMA heritability enrichment analysis of six traits across the three developmental cell stages. SCZ= schizophrenia, BP=bipolar disorder, EA= educational attainment, MDD=major depressive disorder, ASD=autism spectrum disorder, ADHD=attention deficit hyperactivity disorder. **b**, Magma heritability enrichment in random expression matched gene lists from 100 permutations (in blue) compared to the up-regulated genes in neurons (in red).**c,** Expression of the differentially expressed genes contributing to the MAGMA schizophrenia signal (FDR <5%).

**Extended Data Fig. 7.**
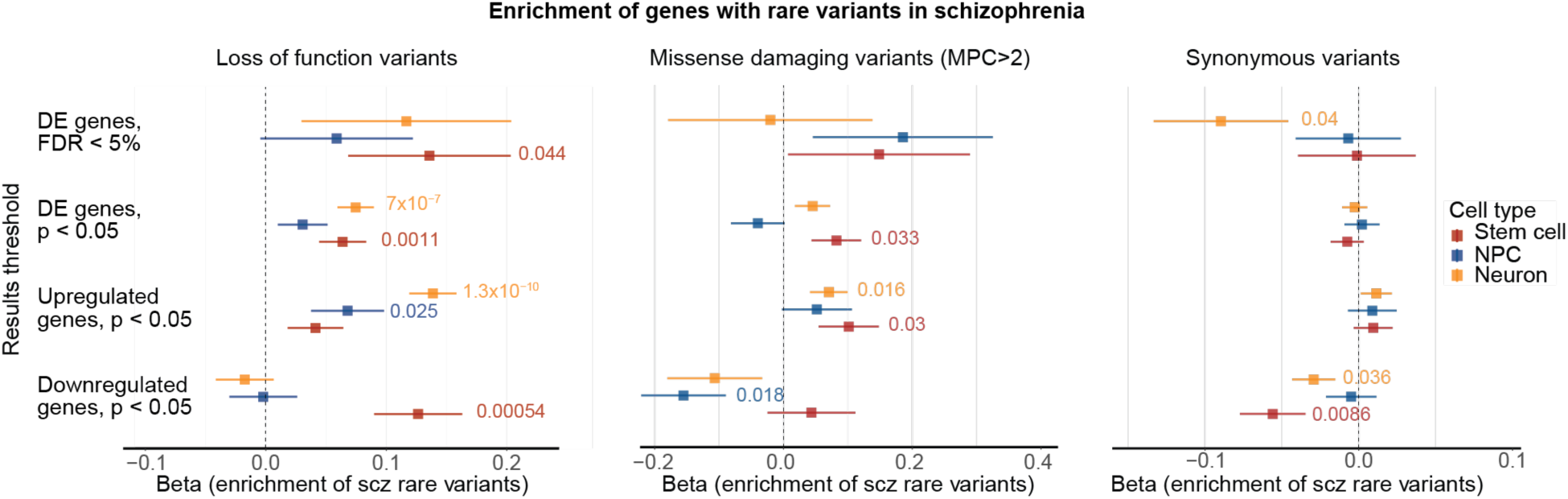
Enrichment analysis of genes with rare variants in schizophrenia. Forest plots for the difference in number of rare coding variants (beta) between schizophrenia patients and controls for loss of function, missense damaging and synonymous variants in genes at different significance cutoffs: FDR <5%, p <0.05 (all genes), upregulated genes with p <0.05 and downregulated genes with p <0.05 at all developmental stages.

**Extended Data Fig. 8.**
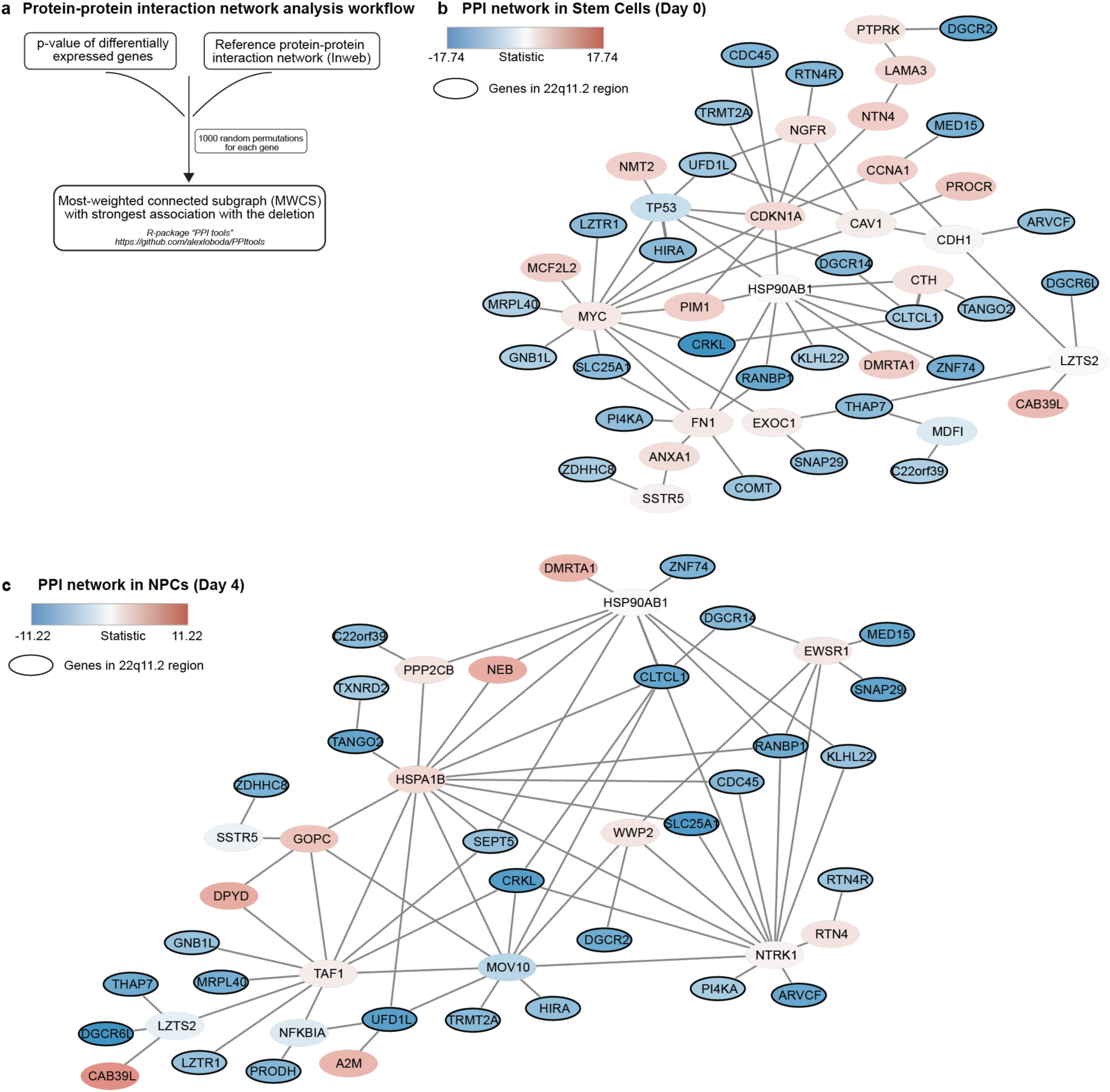
The most weighted sub-cluster graph for protein-protein interactions (PPI) for differentially expressed genes. **a**, Workflow. **b**, Network in Stem cells. **c**, Network in neuronal progenitor cells.

**Extended Data Fig. 9.**
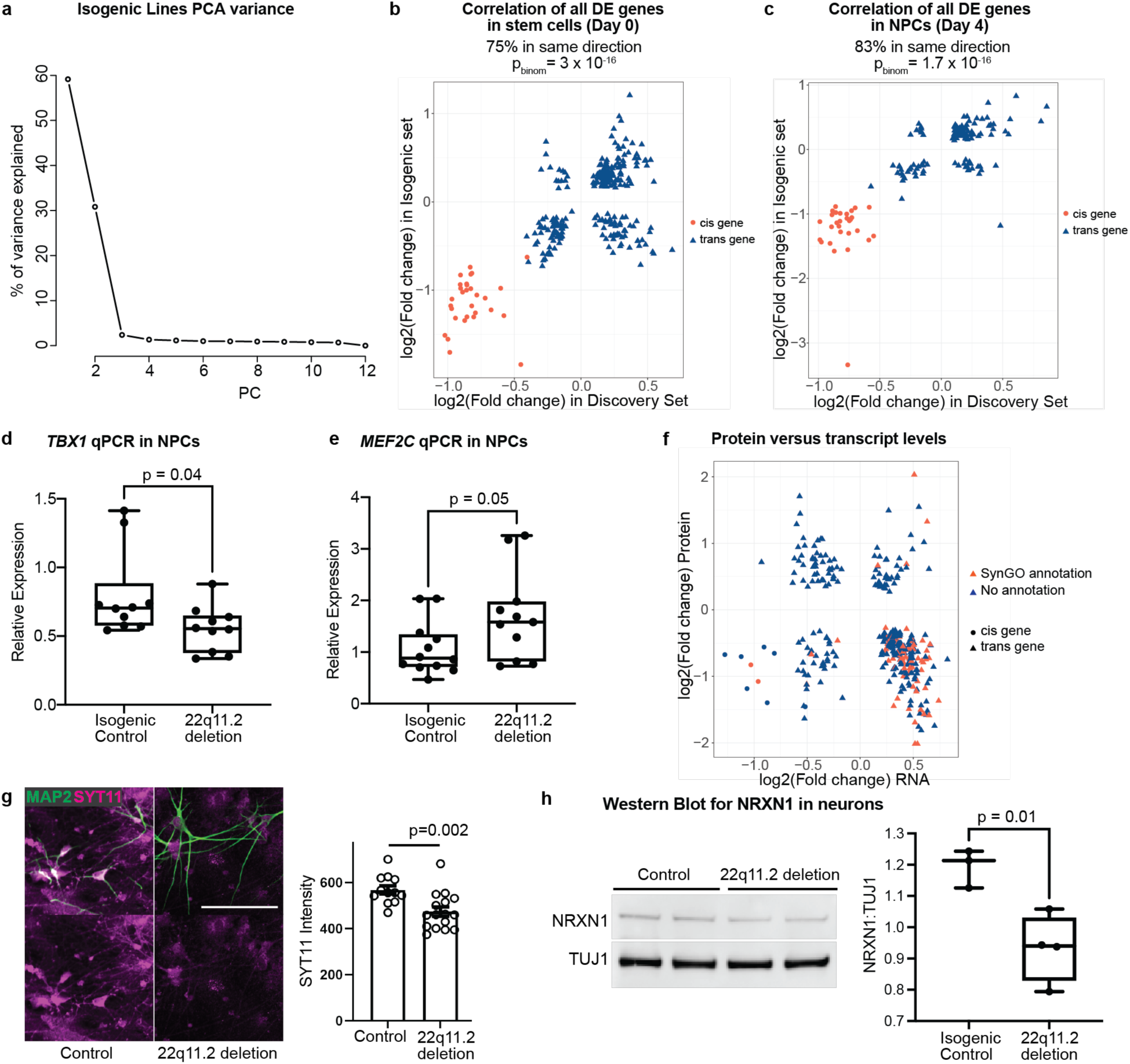
Isogenic line and protein analysis. **a**, Variance explained by each principal component from RNA sequence data in the isogenic lines. **b-c**, Correlation of fold-changes of differentially expressed genes in discovery and isogenic datasets in stem cells (**b**) and neuronal progenitors (**c**). Red circles = cis genes. Blue triangles = trans genes. **d**, Relative expression of TBX1 via qPCR in Day 4 NPCs from isogenic control and 22q11.2 deletion lines (Samples: 2/2, 3BR/2TR, p=0.04). **e**, Relative expression of MEF2C via qPCR in Day 4 NPCs from isogenic control and 22q11.2 deletion lines (Samples: 2/2, 3BR/2TR, p=0.05). **f**, Protein versus RNA levels of genes differentially expressed in 22q11.2 deletion carrier patient neurons. Genes with a SynGO annotation are shown in red, genes with no SynGO annotation are shown in blue. Circles = cis genes. Triangles = trans genes. **g**, Synaptotagmin-11 (SYT11) protein levels are decreased in Day 28 22q11.2 deletion neurons. (Left) Representative confocal images of control and 22q11.2 deletion patient neurons stained with antibodies against SYT11 (magenta) and MAP2 (green). Scale bar is 100 µm. (Right) Quantification of total SYT11 fluorescence within MAP2-positive area normalized to controls. Data are represented as means ± SEM. Individual points are analyzed fields of view from 3 independent control lines and 4 patient-derived lines. Statistical analysis by Student’s t test reveals statistically significant (p=0.0022) decrease in SYT11 levels in patient-derived neurons. **h**, Expression of Neurexin-1 (NRXN1) in total protein lysates from isogenic control and 22q11.2 deletion neurons. (Left) Total protein lysates from isogenic control (left two lanes) and deletion lines (right two lanes) stained for NRXN1 (top) and TUJ1 (bottom). (Right) Statistical analysis by Student’s t test reveals statistically significant decrease in Neurexin-1 expression in the deletion lines. (Samples: 5/5, 2BR/2TR, p=0.01). BR = biological replicate (independent differentiations); TR = technical replicate (independent wells).

## Methods

### Human pluripotent stem cell (hPSC) lines cohort and derivation

We assembled a scaled discovery sample set through highly collaborative, multi-institutional efforts with the Stanley Center Biobank (Broad Institute), the Swedish Schizophrenia Cohort (Karolinska Institute), the Northern Finnish Intellectual Disability Cohort (NFID), Umea University, Massachusetts General Hospital (MGH), McLean Hospital, and GTEx. Human induced pluripotent stem cell (hiPSC) lines were generated from either fibroblasts or lymphoblasts, and either reprogrammed in house (as previously described^34^), at the New York Stem Cell Foundation (NYSCF) or at the Harvard Stem Cell Institute (HSCI) iPS core as listed in Extended Data Table 1. The human embryonic stem cell (hESC) line H1 was obtained from the Human Embryonic Stem Cell Facility of the Harvard Stem Cell Institute.

### hPSC culture

Human ESCs and iPSCs were maintained on plates coated with geltrex (life technologies, A1413301) in StemFlex media (Gibco, A3349401) and passaged with accutase (Gibco, A11105). All cell cultures were maintained at 37°C, 5% CO2.

### Infection of hPSCs with lentiviruses

Lentivirus particles were produced by Alstem (http://www.alstembio.com/). hPSCs were seeded in a geltrex coated 12 well plate at a density of 100,000 cells/cm^2^ in StemFlex medium supplemented with rock inhibitor (Y27632, Stemgent 04-0012) and lentiviruses, at a MOI (multiplicity of infection) of 2. 24 hours later, the medium was changed to StemFlex. The cells were grown until confluency, and then either maintained as stem cells, passaged, banked, or induced with Doxycycline for neuronal differentiation.

### Neuronal differentiation

hPSCs were differentiated into cortical glutamatergic neurons as previously described^34^. Our protocol differs from previous Ngn2-driven protocols^33,89^ through inclusion of developmental patterning alongside Ngn2 programming^34^ (Fig.1b,c,f). This paradigm generates post-mitotic excitatory cortical neurons that are highly homogeneous in terms of cell type^34^ compared to most differentiation paradigms which yield heterogeneous cell types^90^. At 4 days post induction, cells are co-cultured with mouse glia to promote neuronal maturation and synaptic connectivity^91,92^.

### RNA sequencing and alignment

We used triplicate wells of each line at each time point to reduce experimental variation. Cells were harvested in RTLplus Lysis buffer (Qiagen 1053393) and stored at -80°C. To minimize technical biases in readouts from cases and controls, we carried out the RNA sequencing in mixed pools of both genotypes. Sequencing libraries were generated from 100 ng of total RNA using the TruSeq RNA Sample Preparation kit (Illumina RS-122-2303) and quantified using the Qubit fluorometer (Life Technologies) following the manufacturer’s instructions. Libraries were then pooled and sequenced by high output run on a HiSeq 2500 (Illumina). The total population RNA-seq fastq data was aligned against ENSEMBL human reference genome (build GRCh37.p13/hg19) using STAR (v.2.5)^93^. Prior to genome aligning, we used Trimmomatic (v.0.36)^94^ to clip Illumina adapters and low-quality base-pairs from the ends of the sequence reads and removed reads with length < 36 base-pairs. The gene-wise read-counts were generated from the aligned reads by featureCounts in Rsubread (v.1.32)^95^ using GENCODE GTF annotation version 19. The reads from the three experimental replicates were summed together. The final read counts did not differ between cases and controls (11.0 x 10^6^ and 10.8 x 10^6^ reads, respectively; p=0.68, two-sided t-test). The deleted cis genes accounted for 0.53 to 0.71‰ and 0.97 to 1.29 ‰ of all read counts in carriers and controls, respectively.

The plot in Figure 2a was generated as follow: we used normalized read counts from DeSeq2 for a set of 18 canonical marker genes for pluripotency (SOX2, POU5F1, NANOG, and MKI67), neuronal progenitor cells (NEUROD1, SOX2, EMX2, OTX2, HES1, MSI1, and MKI67), and neuronal marker genes (RBFOX3, SYN1, DCX, MAP2, TUBB3, NCAM1, and MAPT) to address the progress of neuronal differentiation in the data set. The normalized gene-wise read counts were scaled to a standard score 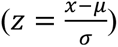 so that the gene expression of the different genes was presented as a difference from the average in units of standard deviations. The mean z-score for each gene set was then calculated and plotted as a line plot across the three cell stages (stem cells, NPCs, and neurons) with 95%-confidence intervals using inbuilt statistics in ggplot2.

### Differential gene expression analysis

For differential gene expression analysis, we applied Wald’s test for read counts that were normalized for library size internally in DESeq2^96^. The differential expression analysis was conducted separately for each cell stage to avoid any biases in gene variance modeling resulting from gene expression differences in between SCs, NPCs, and neurons. The experimental batch was included to the design formula in DESeq2 to correct for the 6 experimental batches in which the data was generated in. We used SVA package (version 3.32)^97^ in R to search for latent factors to remove any unwanted variation in the data. We first estimated the number of latent factors using the leek method in num.sv function that was then used for calculating surrogate variables with irw method and five iterations in sva function. The design model for sva included experimental batch and deletion genotype. One latent factor was identified for the neuron data and was included to the design formula in DESeq2 for differential expression. For Stem cells and NPCs no latent factors were identified. The results for differential expression were obtained for FDR adjusted p-value of < 0.05. A principal component analysis was performed for all genes with more than 10 reads after normalizing the read counts by variance stabilizing transformation in DESeq2. For differential expression analysis in the edited isogenic deletion cell lines we used Limma-voom package^98,99^ that enabled to model the non-independent experimental replicates from each clone with the “duplicateCorrelation” function, which was included in the design model by the block design in Limma.

### Power analysis

The power estimates were calculated using RNASeqPower^100^ (R package version 1.18.0). We calculated the median expression and variance in carriers and controls for all genes with one or more reads (25,264 genes) in the pilot data sets. We assumed equal number of cases and controls, while the coefficient of variance was calculated separately for cases and controls. The alpha level was set to nominal significance of 0.05. For the final data set the power to detect fold changes of >2 was calculated for each gene separately.

### Enrichment for neurodevelopmental and constraint genes

Gene lists for neurodevelopmental disorder genes were compiled from the deciphering developmental delay project^45,46^, and recent large scale exome sequencing study in autism^27^. We included genes for which there was statistical overrepresentation of loss of function variants in patients compared to controls (total 97 genes for ASD ^27^ and 93 for ID^46^ genes). From the earlier DDD-study^45^ we included all “confirmed” developmental disorder genes that affect the brain. We included only those that had “hemizygous” and “monoallelic” as the allelic requirement, and mutation consequence defined as: “loss of function”, “cis-regulatory” or “promotor mutation”, and “increased gene dosage” (total 158 genes). This resulted in a list of total 295 disease genes for neurodevelopmental disorders (Table S5). P-values for the enrichment analyses were calculated with hypergeometric test and binomial test in R. GO-term overrepresentations were calculated with hypergeometric test implemented in GoStats v. 1.7.4^101^ in R with gene identifiers from org.Hs.eg.db. All p-values were calculated for overrepresentation using all mapped genes from each experiment as the background gene universe. False discovery rate (fdr) was used to adjust the raw p-values from the hypergeometric test for overrepresentation using p.adjust function in R. Significance threshold for overrepresentation was set to fdr-adjusted p-value smaller or equal to 0.05. The overrepresentation of synaptic GO terms was estimated by Fisher exact test in the SYNGO online portal (www.syngoportal.org) using a custom background gene set from the RNASeq data set.

### Protein-protein interaction network analysis

Previous efforts have shown that the observed distribution of the p-values from differential expression studies could be modeled as a mixture of the distributed signal and uniformly distributed noise components^102,103^. In such approach, a threshold value could be estimated for observed p-values to discriminate between the likely true signal from noise. Hence, genes could be scored with logarithm of signal to noise ratio (log for making scores additive). Further, using a reference functional network we can leverage gene weights on the map of functional interactions to construct a node-weighted graph. Within this graph a search for the most-weighted connected subgraph (MWCS) could be performed. This search returns a functional module that has the strongest cumulative association to a trait being investigated. Appearance of genes in MWCS is driven both by their differential expression p-value and reference network topology. Thus, non-randomness of each gene’s appearance could be evaluated by randomly permuting p-values and creating a random reference network with preserved node degrees. Estimates of how often a gene will be observed in MWCS by chance provide an empiric metric of significance and could be used to prioritize genes within MWCS. We implemented this strategy in R-package “PPItools” which provides a set of functions to identify MWCS, describe its statistical properties and prioritize genes within it. We used the InWebIM^52^ direct protein-protein interactions network as a reference.

For every time point a beta-uniform mixture distribution was fitted to a distribution of observed p-values. Bonferroni adjusted significance threshold (0.05 / #Genes expressed) was selected as a threshold to discriminate positively and negatively scoring genes. Scores were estimated as a ratio between values of probability density function of Beta distribution at given p-value and threshold p-value or (α-1)×(log(x)-log(x_threshold )), where α is an estimated parameter of Beta distribution. MWCSs for every time point of the experiment (iPSC, neuronal progenitors and neuronal cells) were identified (Fig. 4g and Extended Data Fig. 8). Using described above permutational scheme, for every module we assessed a non-randomness of presence for every gene found in the module (Table S8). After multiple hypothesis testing correction (Bonferroni method used) several genes from each data set come up as significantly functionally enriched (adjusted p < 0.05). 36 out of 50 genes in the iPSC module were seen in random MWCS with less than 5/1000 frequency.

We further tested for excessive connectivity between significantly differentially expressed genes and known neurodevelopmental disease genes. We selected 295 likely disease-causing genes from the Deciphering developmental delay (DDD) project, and a recent, large exome-sequencing study in autism (Table S5). Curated inflammatory bowel disease (IBD) and Parkinson’s disease (PD) risk gene lists (Table S5) were included as a negative control set in this analysis. We estimated the number of connections between genes found in each of the disease gene lists and a list of differentially expressed genes with FDR < 5% normalized to the total number of connections observed for all genes in both tested sets (disease and expression) in reference data. The obtained result could be interpreted as a proportion of all connections that are linking disease and differentially expressed genes. To evaluate significance, we generated random gene sets of the same size as the disease gene sets and estimated an expected number of connections with each set of differentially expressed genes. It is important to note that genes co-expressed within the same tissue or cell type tend to have a greater number of connections between them than would be expected for a random pair of genes. Hence, in generating random gene sets we specifically selected genes at random to match the expression pattern of a disease gene set in a given cell type (iPSC, neuronal progenitors or neurons). For every dataset, the expression distribution was binned into deciles and every gene was assigned to an appropriate bin using mean counts. Random gene sets were selected to match the distribution of genes into deciles for disease gene sets. Empirical p-values were adjusted for two disease gene sets tested with Bonferroni correction.

The PPItools package for finding MWCS and performing network prioritizations along with documentation and source code to perform described analysis is available through GitHub https://github.com/alexloboda/PPItools.

### SNP heritability analysis

LD Score regression^104^ and MAGMA^60^ were used for evaluating common variant associations in and near differentially expressed genes. Briefly for LD score regression, it can be shown that under a basic polygenic model we expect the GWAS statistics for SNP *j* to be:

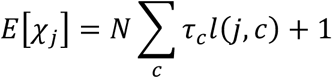

where *N* is the sample size, *c* is the index for the annotation category, lj,c is the LD score of SNP *j* with respect to category *C_c_*, and c is the average per-SNP contribution to heritability of category *C_c_*. That is, the 2 statistic of SNP *j* is expected to be a function of the total sample *N*, how much the SNP tags each category *C_c_* (quantified by lj,c, the sum of the squared correlation coefficient of SNP *j* with each other SNP in a 1 cM window that is annotated as part of category *C_c_*) and c, the effect size of the tagged SNPs.

With this model, LD Score regression allows estimation of each c. Each c is the contribution of category Cc after controlling for all other categories in the model (we included 74 annotations that capture different genomic properties including conservation, epigenetic markers, coding regions and LD structure similar to^105^ and can be interpreted similarly to a coefficient from a linear regression. Testing for significance of c is useful because it indicates whether the per-SNP contribution to heritability of category *C* is significant after accounting for all the other annotations in the model. In addition to considering the conditional contribution of category *C_c_* with c, the total marginal heritability explained by SNPs in category *C_c_*, denoted hg2(Cc), is given by

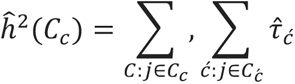

In other words, the heritability in category *C_c_* is the sum of the average per-SNP heritability for all SNPs in *C_c_*, including contributions to per-SNP heritability from other annotations c’ that overlap with category *C_c_* (as indicated by terms of the inner sum where c’≠c). Importantly, *ĥ_g_*(*C_c_*) does not depend on the categories chosen to be in the model and provides an easier interpretation. Therefore, this quantity is the main focus of the analysis.

Here we focus on *ĥ_g_*(*C_c_*) where C_c_ comprises HapMap SNPs 100 kb upstream and downstream of each gene differentially expressed gene. *ĥ_g_*(*C_c_*) was calculated for three sets of differentially expressed genes using two p-value thresholds (FDR < 5% and p < 0.05). Genes surpassing p <0.05 cut-off were further divided to up and down-regulated genes. Heritability estimates were calculated for 6 sets of summary statistics from large GWAS of educational attainment^54^ and 5 psychiatric/neurodevelopmental disorders: ADHD^55^, autism spectrum disorder^56^, bipolar disorder^57^, major depressive disorder^58^ and schizophrenia^59^ OR^32^. In addition, the *ĥ_g_*(*C_c_*) was calculated for the up-regulated genes in neurons (p-value <0.05) and summary statistics for 650 phenotypes from the UK-biobank that have a significant heritability, defined by having a heritability p-value < 0.05 after Bonferroni correction for multiple testing (https://www.nealelab.is/uk-biobank/).

Similar to what was done for LD-score regression we considered gene-lists of differentially expressed genes to ask whether the differentially expressed genes are more strongly associated with each of the six phenotypes. We then used competitive gene set enrichment analysis using gene-wise p-values ^56^ that were calculated for each trait in MAGMA v 1.06 with standard settings^60^. All the results are adjusted for a set of baseline set of covariates with the goal to minimize bias due to gene-specific characteristics: gene size, log(gene size), SNP density, log(SNP density), inverse of the minor allele count, log(inverse of minor allele count) and number of exons in the gene. Gene-wise p-values were calculated by gene analysis in MAGMA and were used to identify genes underlying the stronger association signal among the upregulated genes in neurons. LD-score regression and MAGMA competitive gene set enrichment analyses were repeated for schizophrenia with 100 random genes lists that were matched with expression (±10%) to that of genes that were upregulated in deletion carriers in neurons.

### Analysis of enrichment of differentially expressed genes in whole-exome sequencing data

We investigated if up- and down-regulated genes in 22q11.2 deletion carriers are significantly disrupted by ultra-rare coding variants (URVs) in the whole-exomes of schizophrenia cases and controls (previously described^63,64^). In the cohorts separately, we regressed case status on the number of damaging URVs in the gene set of interest while controlling for the total number of URVs, sex, and the first five principal components. We define damaging URVs as putatively protein-truncating variants (stop-gain, frameshift, and splice-donor and acceptor variants), and damaging missense variants as variants with a MPC score of >= 2, as previously described^106^. We applied inverse-weighted meta-analysis to combine the test-statistics from both studies to get a single joint P-value. We tested for enrichment in up- and down-regulated genes, and a collection of randomly sampled neuronally-expressed genes.

### Motif enrichment analysis

The motif enrichment analysis was carried out by Homer software for genes whose transcripts were found upregulated (log_2_ Fold change>0) at day 28 neurons and p-value below < 0.05. We performed a *de novo* motif analysis for human motifs using findMotifs.pl with len = 10. We curated the obtained results by setting a stringent p-value threshold (p < 10^-10^), visually inspecting that observed motifs do not match only from the edges, excluded repeat sequences, and required that the motif had a frequency of above 5%.

### CRISPR generation of isogenic 22q11.2 cell lines

To generate an isogenic 22q11.2 line in H1 hESCs, oligonucleotides (IDT) targeting LCR A (ACACTGGGCACATTATAGGG) and LCR D (CATTCATCTGTCCACCCACG) were cloned into a pU6-sgRNA vector generate sgRNA plasmids pPN298 and pPN306, respectively, via procedures described previously ^107^. For transfection, cells were pre-incubated with “1:1 medium” composed of a 1:1 mixture of mTeSR1 medium and “hPSC medium” [hPSC medium: KO DMEM (Gibco 10829-018) with 20% KOSR (Gibco 10828-028), 1% Glutamax (Gibco 35050-061), 1% NEAA (Corning 25-025-Cl), 0.1% 2-mercaptoethanol (Gibco 21985-023) and 20ng/ml bFGF (EMD Millipore GF0003AF) supplemented with 10μM ROCK inhibitor (Υ-27632). 7 μg Cas9 nuclease plasmid (pX459, Addgene #62988) 1.4 μg pPN298 and 1.4 μg pPN306 were electroporated into 2.5×106 cells at 1050V, 30ms, 2 pulses (NEON, Life Technologies MPK10096), as described ^108^. Individual hPSC colonies were selected with puromycin treatment and seeded into Geltrex-coated 96-well plates, expanded for 1-2 weeks and duplicated for cell freezing and gDNA extraction. Clones were frozen in 96-well plates using 50% 1:1 medium plus 10μM Υ-27632, 40% ¬FBS (VWR SH30070.03) and 10% DMSO (Sigma D2650). gDNA was extracted overnight at 55°C in Tail Lysis Buffer (Viagen 102-T) with Proteinase K (Roche 03115828001) followed by a 1hr 90°C incubation. Droplet digital PCR (ddPCR) was performed to determine for copy numbers of the HIRA and ZNF74 genes using probes previously described^109^. SNP genotyping was performed using the Illumina Infinium PsychArray-24 Kit on the lines to confirm the microdeletion (Broad Institute, Cambridge, MA). Differential expression for the isogenic lines was performed by DESeq2. The results from isogenic lines were compared to the results obtained from the discovery sample. The overlap between the direction of fold-changes in isogenic samples were tested using binomial test for all genes that were differentially expressed in the discovery sample. The expected probability for overlap was calculated from all genes and was on average 0.5. The differences in gene expression were tested by Mann-Whitney test including all genes with nominally significant p-value in differential expression in the isogenic lines.

### DNA FISH analysis

FISH (Fluorescent In-Situ Hybridization) analysis was conducted in the isogenic control and 22q11.2 deletion lines to analyze the copy number of the 22q11.2 region and validate the isogenic deletion. We generated the probe using a bacterial artificial chromosome (BAC) located in the 22q11.2 region, CTD-2300P14 (Thermo Fisher Scientific, Supplier Item: 96012), labeled with Cy3 dUTPs (GE healthcare: PA53022), by means of nick translation (Abbott: 32-801300), and visualized the labeled cells using confocal microscopy.

### Multielectrode Arrays (MEA)

MEA experiments and analysis were performed exactly as previously described^34^. Briefly, neuronal progenitors (at day 4) were seeded on 8×8 MEA grids, each with 64 microelectrodes, in the absence or presence of mouse glia, and routinely sampled these for 42 days after Ngn2 induction and dual SMAD and WNT inhibition. Each MEA plate contained wells from both deletion carrier and control neurons to minimize technical biases. Extracellular spikes (action potentials) were acquired using Axion Biosystems multi-well MEA plate system (The Maestro, Axion Biosystems; 64 electrodes per culture well). During the recording period, the plate temperature was maintained at 37± 0.1 °C, environmental gas composition was not maintained outside of the incubator. Unless otherwise stated, descriptive statistics for MEA data is presented as Tukey style box plots, showing the 1st, 2nd, and 3rd quantile (Q1, Q2, & Q3 respectively; inter-quartile range, IQR = Q3-Q1). Box plot whiskers extend to the most extreme data points between Q1-1.5*IQR and Q3+1.5*IQR ^110–112^. All data points outside the whiskers are plotted. Non-parametric 95 % confidence intervals for M are calculated using fractional order statistics ^113^.

### TMT-processing workflow

Cell pellets were lysed and 50ug protein per TMT channel were subjected to disulfide bond reduction and alkylation. Methanol-chloroform precipitation was performed prior to protease digestion with LysC/trypsin. Obtained peptides were labeled with the respective TMT reagents and pooled. Enhanced proteome coverage was achieved by high-pH reversed phase fractionation to reduce sample complexity. Peptide fractions were analyzed on an Orbitrap Fusion mass spectrometer using SPS-MS^114^. Mass spectra were processed using a Sequest-based in-house software pipeline. Peptide and protein identifications were obtained following database searching against all entries from the human UniProt database. For TMT-based reporter ion quantitation, we extracted the summed signal-to-noise (S:N) ratio for each TMT channel. For protein-level comparisons, peptide-spectrum-matches (PSM) were identified, quantified, and collapsed to a 1% peptide false discovery rate (FDR) and then collapsed further to a final protein-level FDR of 1%. Moreover, protein assembly was guided by principles of parsimony to produce the smallest set of proteins necessary to account for all observed peptides. Proteins were quantified by summing reporter ion counts across all matching PSMs using in-house software. Protein quantification values were exported for further analysis.

### Analysis of protein abundances

Differences in protein abundances between deletion carriers and controls were estimated in day 28 neurons derived from two patient (SCBB1962 and SCBB-1825) and two control lines (SCBB1828, SCBB1827) in total 18 replicates. The abundances for the detected 8811 gene products were log_2_+1 transformed and quantile normalized in Limma package^99^ (v. 3.3.49) in R. A linear model including instrument run and deletion status was used to analyze differences in the normalized protein abundances between deletion carriers and controls in Limma. The correlation of the non-independent experimental replicates was estimated with “duplicateCorrelation” function (average estimated inter replicate correlation was 0.83) and was taken into account in the design model using block design in Limma. Overlap of gene products between RNA sequence data and proteomics data (total 8585 gene products detected by both methods) was compared using p-value<0.05 threshold. The overlap of direction of effect was estimated with binomial test with expected probability of 0.5. The density coloring was calculated from Kernel density estimation using densCols in R.

### Immunohistochemistry

Cultured induced neurons were fixed in 4% paraformaldehyde + 20% sucrose in DPBS for 20 min at room temperature. Cells were incubated with blocking buffer containing 4% horse serum, 0.1M Glycine, and 0.3% Triton-X in PBS for 1 hour at room temperature. Primary antibodies, diluted in 4% horse serum in PBS, were incubated overnight at 4oC. Secondary antibodies were diluted in 4% horse serum and applied for 1 hour at room temperature. Samples were washed 3x with PBS and imaged on spinning disc confocal microscope (Andor Dragonfly) with a 20x air objective. The following antibodies were used: rabbit anti-SV2A (1:1000, Abcam ab32942), chicken anti-MAP2 (1:10,000, Abcam ab5392), rabbit anti-Synaptotagmin-11 (1;1000, Synaptic Systems 270 003). Alexafluor plus-555 and Alexafluor plus-488 conjugated secondary antibodies (1:5,000) were obtained from Invitrogen.

### Image acquisition and analysis

Fluorescent images were acquired on spinning disc confocal microscope (Andor Dragonfly) at room temperature using 20x air interface objective using Fusion software. For quantification at least four 1024×1024 pixel fields of view from 2 different wells were taken for each line. The images were analyzed using ImageJ software.

### Immunoblotting

For collection, neurons grown on glia were washed with DPBS and lysed with RIPA buffer and 1x protease inhibitor cocktail. Lysates were boiled, sonicated and centrifuged at 16,000xg for 5 minutes. The soluble fraction was separated on SDS-PAGE using Bolt system (Novex). The proteins were transferred onto nitrocellulose membrane using iBlot2 Gel Transfer Device and immunostained using Neurexin-1 antibody (Millipore ABN161-I) and Tuj1 (Biolegend 801201) and detected via HRP-conjugated secondary antibodies on the Chemidoc system.

### qPCR analysis

RNA isolation was performed with the Direct-Zol RNA miniprep kit (ZYMO: cat# R2051) according to the manufacturer’s instructions. To prevent DNA contamination, RNA was treated with DNase I (ZYMO: cat# R2051). The yield of RNA was determined with a Denovix DS-11 Series Spectrophotometer (Denovix). 200ng of RNA was reverse-transcribed with the iScript cDNA Synthesis Kit (Bio-Rad, cat# 1708890). For all analyses, RT–qPCR was carried out with iQ SYBR Green Supermix (Bio-Rad, cat# 1708880) and specific primers for each gene (Supplementary Table) with a CFX384 Touch Real-Time PCR Detection System (Bio-Rad). Target genes were normalized to the geometric mean of control genes, RPL10 and GAPDH, and relative expression compared to the mean Ct values for control and wild-type isogenic samples, respectively.

The following primers were used:

MEF2C_forward 5’-CTGGTGTAACACATCGACCTC-3’
MEF2C_reverse 5’-GATTGCCATACCCGTTCCCT-3’
TBX1_forward 5’-ACGACAACGGCCACATTATTC-3’
TBX1_reverse 5’-CCTCGGCATATTTCTCGCTATCT-3’
RPL10_forward 5’-GCCGTACCCAAAGTCTCGC-3’
RPL10_reverse 5’-CACAAAGCGGAAACTCATCCA-3’
GAPDH_forward 5’-GGAGCGAGATCCCTCCAAAAT-3’
GAPDH_reverse 5’-GGCTGTTGTCATACTTCTCATGG-3’

